# Tumor reactivity assessment using clonal expression (TRACE) reveals tumor reactive CD8^+^ T cell heterogeneity across solid tumors

**DOI:** 10.64898/2026.02.25.707942

**Authors:** David Monteiro, Jack Denebeim, Anne E. Dodson, Ashish Yeri, Mitali Ghose, Meghan Travers, Sophie Capobianco, Conor Calnan, Gustavo J. Martinez, Charles H. Yoon, Karrie Wong, Micah J. Benson, Dipen Sangurdekar

**Affiliations:** KSQ Therapeutics, Inc., Lexington, MA; Brigham and Women’s Hospital, Harvard Medical School, Boston, MA

**Author notes:** **Correspondence:** Dipen Sangurdekar.

**Keywords:** Tumor infiltrating lymphocytes (TIL), T cell, T cell receptor, tumor antigens, machine learning, biomarkers

## Abstract

1

**Introduction:** Tumor infiltrating lymphocytes (TIL) drive the anti-tumor activity of a broad class of immunotherapies. *In situ* TIL are composed of T cells that recognize tumor antigens (Tumor Reactive T cells, or TRTs) as well as bystander T cells with specificity for other antigens. TRT clonotypes are associated with a unique and tumor-driven exhausted transcriptional state, enabling single-cell RNA sequencing (scRNA-seq)-based predictive models for TRTs using experimentally validated clone labels.

**Methods:** In this study, a clonotype-level CD8^+^ TRT classifier (TRACE) was built using an aggregated dataset of validated tumor reactive clonotypes and associated scRNA-seq data from multiple publications that overcomes the limitations of training on a single dataset, donor, or indication. TRACE does not require dataset manipulation for training or prediction, enabling it to be easily applied to new test datasets as they emerge.

**Results:** TRACE exhibited robust performance on held-out TIL and PBMC clones - achieving a mean Matthews correlation coefficient of 0.84 and F1-score of 0.85 - comparable to or outperforming other TRT prediction methods. We experimentally confirmed the reactivity of TRACE-identified TRT clones by co-culturing engineered, *ex vivo* expanded TIL with autologous melanoma tumor cell lines. Finally, we applied TRACE to evaluate the frequency of TRT across hundreds of patient samples from multiple tumor atlases spanning lung, colorectal, and pancreatic cancer. TRACE scores were observed to be significantly higher in exhausted CD8 T cells in tumors but not in exhausted cells in normal adjacent or non-cancer samples, suggesting specificity towards identifying tumor-antigen experienced T cells.

**Conclusion:** TRACE is a tumor reactivity scoring algorithm released with open model weights that can be applied to tissue or blood single-cell RNAseq datasets. Its application should be of general interest for characterizing the fraction of TRTs in TIL and for establishing correlations with clinical response to immunotherapies.

## 2 Introduction

Immune checkpoint blockade (ICB) and adoptive cell therapies (ACT), including tumor infiltrating lymphocyte (TIL) therapy, have demonstrated durable clinical benefit across multiple cancers, particularly immunologically hot tumors such as melanoma (1,2). A consistent biological determinant of response to these immunotherapies is the presence of cytotoxic CD8 T cells within the tumor microenvironment (TME), where they mediate direct tumor cell killing and shape local immune dynamics (1,3–5). However, the mere abundance of intratumoral CD8 T cells is insufficient to explain therapeutic efficacy, as tumors often harbor a heterogeneous mixture of tumor reactive and bystander T cells that differ markedly in antigen specificity, transcriptional state, and functional potential (6). Recent studies have established that a substantial fraction of CD8 T cells infiltrating tumors are bystander T cells, recruited independently of tumor antigen recognition and lacking direct anti-tumor activity (6–8). In contrast, increased infiltration of tumor reactive T cells (TRTs) - defined by recognition of tumor-derived antigens, including neoantigens - is preferentially associated with favorable prognosis and response to immunotherapy (9,10). In melanoma and other inflamed tumors, therapeutic response correlates more strongly with the abundance, clonal dynamics, and functional state of TRTs than with total CD8 T cell density (6,8,11–14). These findings underscore the importance of discriminating tumor reactive from non-tumor reactive T cells within complex intratumoral immune populations.

Antigen-experienced TRTs occupy a distinct transcriptional niche within the TME. Multiple single-cell and functional studies have shown that TRTs frequently adopt a tissue-resident memory-like phenotype characterized by expression of markers such as *ENTPD1* (CD39), *ITGAE* (CD103), and chemokines including *CXCL13*, alongside features of chronic antigen stimulation and exhaustion(3,8,10,11,15). Importantly, these transcriptional programs are enriched for tumor reactivity across cancer types and have been linked to both response to checkpoint blockade and persistence following ACT (3,11,16). Nevertheless, no single marker or gene signature fully captures tumor reactivity, and substantial overlap exists between TRTs and non-reactive exhausted or activated T cell states.

This limitation has direct implications for ACT. Clinical and translational studies demonstrate that enrichment of tumor reactive T cells within TIL products using neoantigen screening or phenotypic selection strategies can improve therapeutic efficacy (8,17–21). However, current approaches to identifying TRTs rely on labor-intensive functional assays, peptide-MHC multimers, or bulk phenotypic proxies that are difficult to scale and often incompatible with routine single-cell profiling. Computational approaches applied to single-cell transcriptomic data offer a principled opportunity to address this gap. Recent single-cell studies have substantially advanced the transcriptional characterization of TRTs by integrating scRNA-seq with TCR sequencing and functional validation, revealing conserved programs associated with chronic antigen stimulation, tissue residency, and exhaustion-like states (22–26). Building on these datasets, several computational approaches have attempted to infer tumor reactivity directly from transcriptomic profiles, including marker-based scores, gene signatures, and supervised machine-learning classifiers trained on experimentally validated clonotypes. Recent machine-learning approaches such as TRTpred (26), MANAscore (27), NeoTCR8 (24), and predicTCR (28) represent important steps toward computational identification of tumor reactive T cells by leveraging experimentally annotated single-cell datasets and TCR-linked supervision.

Despite their advances, existing methods share important limitations. First, most models are trained on small, experiment-specific datasets derived from individual studies or tumor types, limiting statistical power and robustness to cohort-specific effects. Second, training data are typically not augmented across experiments or platforms, and model weights are not shared, hindering reproducibility and reuse as new datasets emerge. Third, many approaches require extensive single-cell normalization, batch correction, and hand-tuned feature selection, introducing dataset-specific assumptions and constraining generalizability across sequencing depth, chemistry, and platform. Importantly, while multiple studies demonstrate that cells sharing an identical TCR clonotype can occupy diverse transcriptional states within the TME (3,11,29–31), existing methods (except TRTpred and predicTCR) largely operate at the single-cell level and do not explicitly model intra-clonal heterogeneity or leverage clone structure during learning. Together, these limitations motivate the development of a more scalable, robust, and clonotype-aware framework for inferring tumor reactivity from single-cell transcriptomic data.

To address these challenges, we introduce **TRACE**, a machine-learning framework designed to infer tumor reactivity from scRNA-seq while explicitly accounting for clonal structure and technical variability. In the absence of clonal information, TRACE can be applied to individual cells. When TCR information is available, TRACE incorporates a clone-summarizing strategy in which cells sharing a TCR clonotype are aggregated using optimized parameters that preserve biologically meaningful heterogeneity while reducing noise driven by stochastic gene expression. To further stabilize learning across datasets, TRACE employs expression binning, which discretizes transcript abundance into robust ordinal representations that mitigate sensitivity to sequencing depth, platform differences, and batch effects, while dramatically reducing training time. Model training is guided by systematic hyperparameter optimization, enabling consistent performance across heterogeneous cohorts without manual tuning. Finally, TRACE is released with versioned, openly shared model checkpoints, allowing the community to apply, benchmark, and incrementally update the model as new annotated datasets become available. Collectively, these design choices enable TRACE to overcome key limitations of prior approaches and provide a generalizable, extensible foundation for transcriptome-based identification of tumor reactive T cells.

We performed *in silico* benchmarking of TRACE against other methods with publicly available code or gene signatures and found that TRACE delivers comparable or superior performance across holdout datasets spanning multiple cancer indications. We further experimentally validated TRACE in melanoma by identifying tumor reactive clones through T cell activation, as indicated by 4-1BB expression, in a co-culture assay using *ex vivo* expanded TIL and autologous tumor cells. To demonstrate TRACE’s scalability and utility across diverse datasets, we applied the model to multiple single cell tumor atlases and examined its associations with tumor and clinical metadata. Consistent with its design, TRACE scores were enriched in clonally expanded, exhaustion associated T cells within the tumor but not in adjacent normal tissues. Together, these results highlight TRACE as a robust and generalizable tool for identifying and characterizing tumor reactive T cells across heterogeneous single cell datasets spanning multiple cancer indications.

## 3 Results

### Optimization of TRACE model parameters and training strategy

To build TRACE, we first compiled six scRNA/scTCR-seq datasets spanning multiple indications containing CD8^+^ TIL transcriptomes with matched, experimentally verified tumor reactivity labels (22–26,32) (Table 1). After dataset cleaning, these yielded 9,488 TRTs from 274 clones for training. In addition to experimentally verified non-TRT labels provided by some groups, we supplemented our dataset with a diverse set of probable true negatives. These included published CD8^+^ T cell clones from PBMC samples from healthy (33,34) or COVID-19-experienced (35) individuals in addition to TIL clones with known antigens present in VDJdb (36). In total, 40,156 cells across 16,465 clones were used for model training.

**Table 1.** TRACE was trained on nine scRNA/scTCR-seq datasets (6 TIL + 3 PBMC) containing CD8^+^ T cells with paired TCR_IZβ_ chain information. For each TIL dataset, TRT clones were identified experimentally by the original authors. Non-TRT clones in TIL were either identified as non-tumor reactive by the authors or contained a TCR_β_ with a known antigen in VDJdb. All clones in PBMC samples from healthy or CoV2-experienced patients were considered non-tumor reactive.

| Dataset | Tissue context | Experimentally-validated TRT clones (cells) | Experimentally-validated non-TRT clones (cells) | VDJdb-matched non-TRT clones (cells) | PBMC-derived non-TRT clones (cells) |
| --- | --- | --- | --- | --- | --- |
| Caushi *et al.* (22) | Lung (NSCLC) | 25 (1512) | 0 | 1332 (5833) |  |
| Hanada *et al.* (23) | Lung (NSCLC) | 19 (576) | 0 | 516 (986) |  |
| Lowery *et al.* (24) | Melanoma, breast, GI | 45 (378) | 0 | 400 (707) |  |
| Meng *et al.* (32) | PDAC | 61 (626) | 52 (1510) | 0 |  |
| Oliveira *et al.* (25) | Melanoma | 46 (5006) | 92 (2427) | 10 (164) |  |
| Pétremand *et al.* (26) | Melanoma | 78 (1390) | 96 (1172) | 12 (158) |  |
| Gao *et al.* (33) | Healthy PBMC |  |  |  | 11730 (13317) |
| Ogura *et al.* (35) | Healthy + CoV2-experienced PBMC |  |  |  | 819 (3226) |
| 10x Genomics PBMC (34) | Healthy PBMC |  |  |  | 1132 (1168) |
| **Totals:** |  | 274 (9488) | 240 (5109) | 2270 (7848) | 13681 (17711) |

To eliminate data leakage between training and test sets, we performed train/test splits at the clone-level and rigorously evaluated performance across multiple seeds to select optimal model input parameters (Figure 1, Methods, and Supplementary Data), including 1) the method for collapsing gene expression data to the clone-level, 2) the gene expression transformation method (log normalization or expression binning), and 3) the number and identity of the most important features used in the final model. The final gene list was established by selecting the top *n* ranked genes learned from each training set, defined using median feature importance across all CV folds.

**Figure 1.**
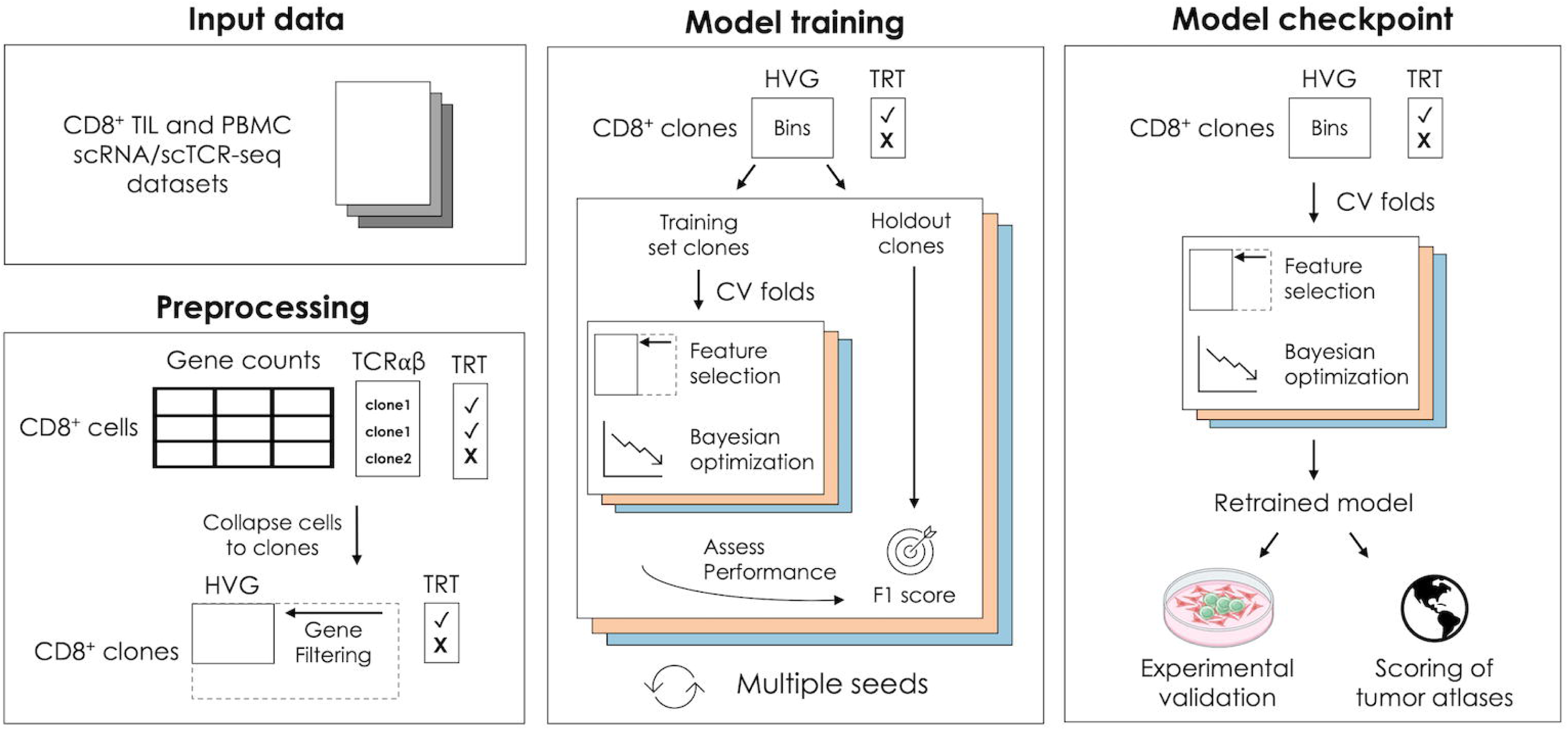
Overview of model training and checkpointing. Single-cell gene expression and paired TCRl1lβ sequences from TIL and PBMC-derived CD8^+^ T cells were used as inputs for model development. Low-expressing genes were filtered out and gene counts for highly variable genes were collapsed to the clone-level and normalized. To build TRACE, CD8^+^ clones were split into training and holdout sets. Cross-validation folds were used to perform feature selection and Bayesian hyperparameter tuning within the training set, and model performance was evaluated on held-out clones. This procedure was repeated across multiple random train/test splits. A TRACE checkpoint was generated after training on the full dataset using optimized hyperparameters.

Across all combinations of these parameters, we observed that model performance, measured by Matthews Correlation Coefficient (MCC) on the holdout test dataset, increased with the number of retained features up to 50 genes, with only a modest increase at higher model sizes (Figure 2A, Supplementary Figure 1). Similar tests identified the optimal method for cell-to-clone gene summarization as the 75^th^ percentile of expression levels across cells within a clone, which matched or outperformed the 50^th^ and 100^th^ percentiles (Supplementary Figure 1). Likewise, we found that expression binning resulted in intermediate models with higher performance across all conditions tested. A comparison of 3 modeling frameworks - XGBoost, AdaBoost, and Random Forest - identified XGBoost as the most performant on this TRT classification task (Supplementary Figure 2).

**Figure 2.**
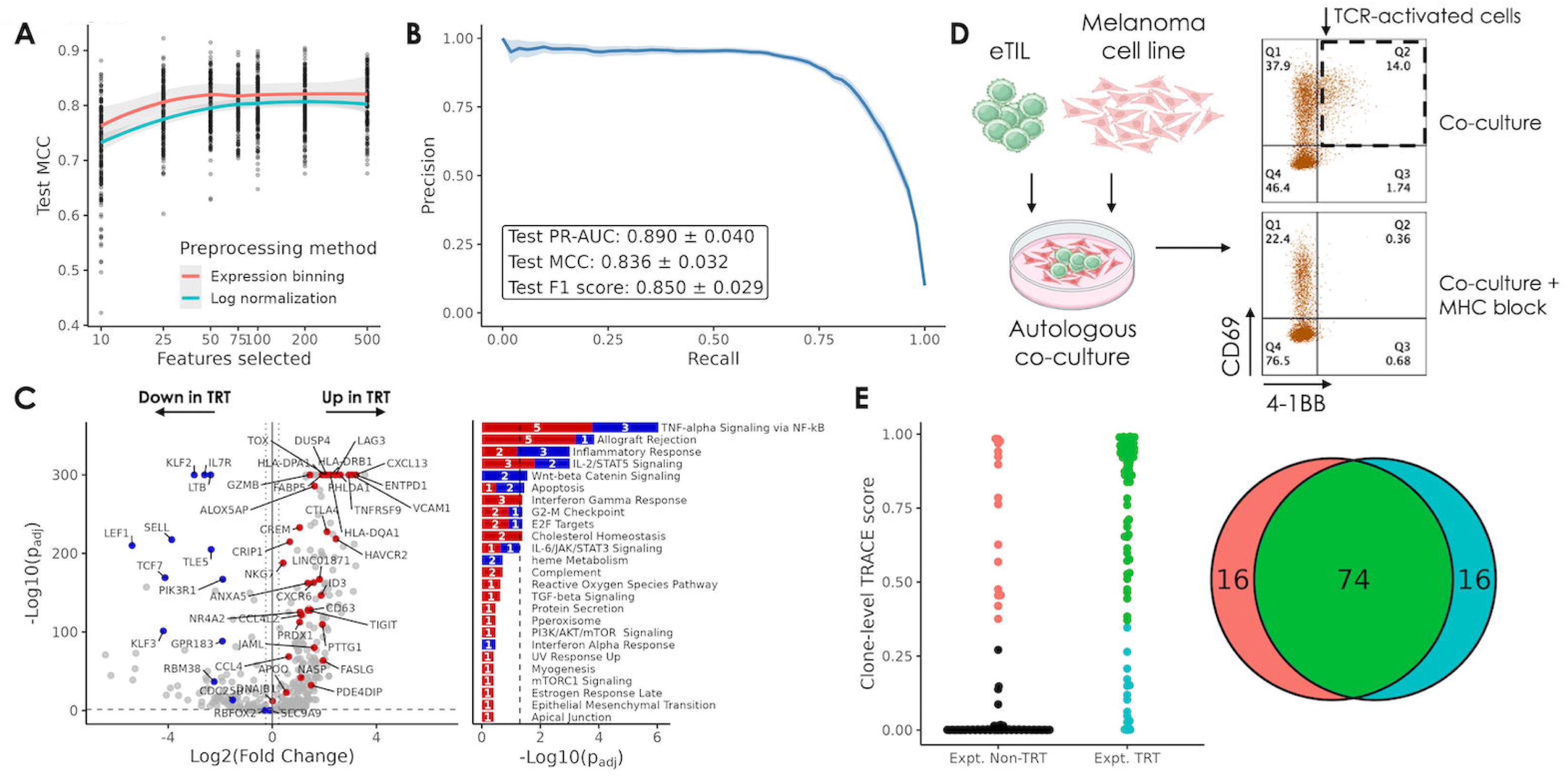
TRACE performance and experimental validation. **(A)** Performance of intermediate TRACE models during model tuning; lines show mean MCC by preprocessing method with a 75^th^ percentile cell-to-clone gene expression collapsing method and Optuna hyperparameter tuning. **(B)** Precision-recall curve (mean with 95% confidence intervals) summarizing TRACE performance across 50 seeds with optimal hyperparameters. Inset shows mean +/- SD. **(C)** Volcano plot highlighting the 50 genes used by TRACE. Grey genes are other highly variable genes used to establish the bins but not used in the final model. Bar plot shows Hallmark pathways containing TRACE genes. On volcano plot, dashed horizontal line represents FDR = 0.05 and dashed vertical lines represent |log_2_(FC)| = 0.25. On bar plot, dashed vertical line represents FDR = 0.05. **(D)** Schematic of autologous co-culture experiments used to validate TRACE performance. **(E)** Summary of overlap between clones predicted to be TRT by TRACE and those exhibiting reactivity during autologous co-culture.

### TRACE accurately identifies validated tumor reactive clones across solid tumors

Using these optimized input parameters (expression binning, 75^th^ percentile, 50 top genes), we performed nested cross-validation across train/test splits from 50 random seeds. Gene selection, hyperparameter tuning, and model training were performed independently on the training data within each iteration, and the trained model was used to predict the tumor reactivity labels of holdout clones. Across these, TRACE exhibited consistently high performance, achieving a mean MCC score of 0.84, mean precision-recall area under the curve (PR-AUC) of 0.89 and mean F1 score of 0.85 (Figure 2B) on holdout test clone data. Model features were selected from one representative iteration, and the final model was retrained with new hyperparameter tuning on all available data to create a TRACE model checkpoint. Notably, the list of 50 gene features was stable across multiple iterations.

Of the 50 genes comprising the TRACE feature set, 36 genes have higher expression on average in TRT clones vs. non-TRT clones in the aggregated dataset, while the remaining 14 have lower expression (Figure 2C). Overall, the up-regulated genes in TRT clones represent a composite phenotype spanning effector-memory, cytotoxicity (*GZMB*, *NKG7*, *CCL4*), and chronic activation/exhaustion (*DUSP4*, *NR4A2*, *ENTPD1*/CD39, *PDCD1*, *LAG3*, *TOX*, *TIGIT*, *CTLA4*). Down-regulated genes included genes expressed in naïve or stem-like T cells (*LEF1*, *IL7R*, *SELL*, *TCF7*, *KLF2*) reflecting the non-antigen experienced or PBMC clones. Gene set and pathway enrichment analysis revealed that among the most-enriched pathways were the TNF-alpha (NF-kB), IFN-gamma, and IL-2/STAT5 pathways, involved in T cell activation and proliferation. Several genes such as *CXCL13*, *ENTPD1*, *TNFRSF9* (4-1BB), and *IL7R*, feature in other published methods (Supplementary Table 3, Supplementary Figure 3).

We established an experimental approach to identify tumor reactive clones in *ex vivo* expanded TIL by co-culturing them with an autologous tumor cell line (Methods). After co-culture, we isolated CD8^+^ T cells for analysis by flow cytometry and confirmed the presence of a population of CD69^+^4-1BB^+^ cells that exhibited MHC-mediated functional reactivity (Figure 2D). For one melanoma donor, we performed scRNA/scTCR-seq on T cells from tumor starting material (TIL) and on T cells from four biological co-culture replicates where we found the same reactive population - with high expression of the gene *TNFRSF9* (encoding 4-1BB). We used these cells to assign experimentally reactive TRT (Expt. TRT) or non-TRT labels to 1,805 clones. Independently, we applied TRACE to tumor starting material and found high concordance between TRACE predictions and Expt. TRT labels. Since TRACE depends on sequencing performed on limited tumor material and Expt. TRT is assessed on *ex vivo* expanded T cells, not all clones with TRACE labels can be validated experimentally using this approach. Among the 171 clones found in both sample types, TRACE correctly predicted 82% (74/90) of Expt. TRT clones and 80% (65/81) of Expt. non-TRT clones (Figure 2E).

### TRACE exhibits comparable or superior performance to other tumor reactivity prediction methods

The TRACE model is trained on multiple validated tumor reactivity datasets, which were also used in the corresponding studies to develop tumor reactivity prediction tools. A thorough methodological comparison of these tools is in Supplementary Table 2. Since TRACE is trained on an aggregated dataset which was further augmented with negative class labels, we expect the model performance of TRACE to be comparable or superior to all other methods. In addition, TRACE code and model checkpoints are publicly available for re-training and prediction on new datasets, whereas not all TRT methods have either code or checkpoints freely available for thorough benchmarking of TRACE. We implemented and retrained four published TRT methods - NeoTCR8, TRTpred, TR30, and MANAscore - and evaluated them against TRACE on our combined dataset (Table 2, Methods). Across the same 50 splits used previously (Figure 2B), TRACE models exhibited the highest mean MCC, comparable to TRTpred and outperforming the TR30, NeoTCR8, and MANAscore methods (Figure 3A). Subsequently, we investigated the performance of these methods on individual published dataset subset from the 50 test splits (Figure 3B). Individual methods tended to perform better on the datasets from which they were developed, including NeoTCR8 (trained on data from Lowery *et al.*) and MANAscore (trained on data from Caushi *et al.* and Oliveira *et al.*) TRACE exhibited the strongest performance in 3 of 4 datasets evaluated and had comparable performance across multiple metrics to the TR30 and TRTpred methods on the Pétremand dataset (Supplementary Table 4). We also assessed the performance of all methods on a combined PBMC subset, where TRACE exhibited a low false positive rate (FPR) in-line with that achieved by TRTpred and MANAscore (Figure 3C). The higher false positive rate of the NeoTCR8 and TR30 methods on this subset could be attributed to the lack of non-TRT-associated genes in these gene sets, which might improve the specificity of TRT classifiers.

**Figure 3.**
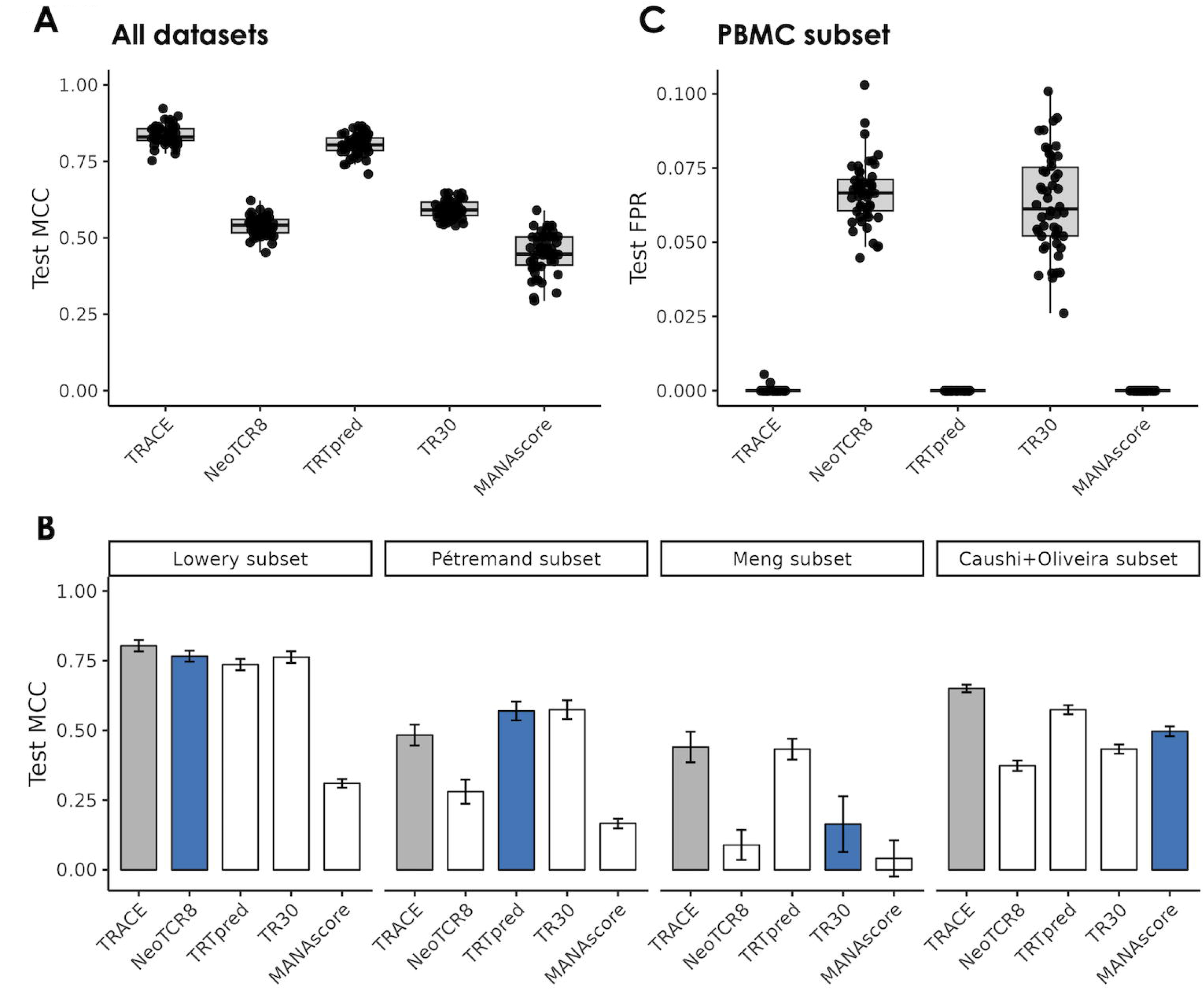
TRACE performance relative to other TRT methods. (A, **B)** Performance (test set MCC) of TRACE and other TRT methods across 50 seeds on test sets containing clones from all available datasets (A) and test sets subset for clones from specific datasets (B). Blue bars represent methods trained on the respective datasets. **(C)** False positive rate of all methods across 50 test sets subset for clones from PBMC datasets.

**Table 2.**
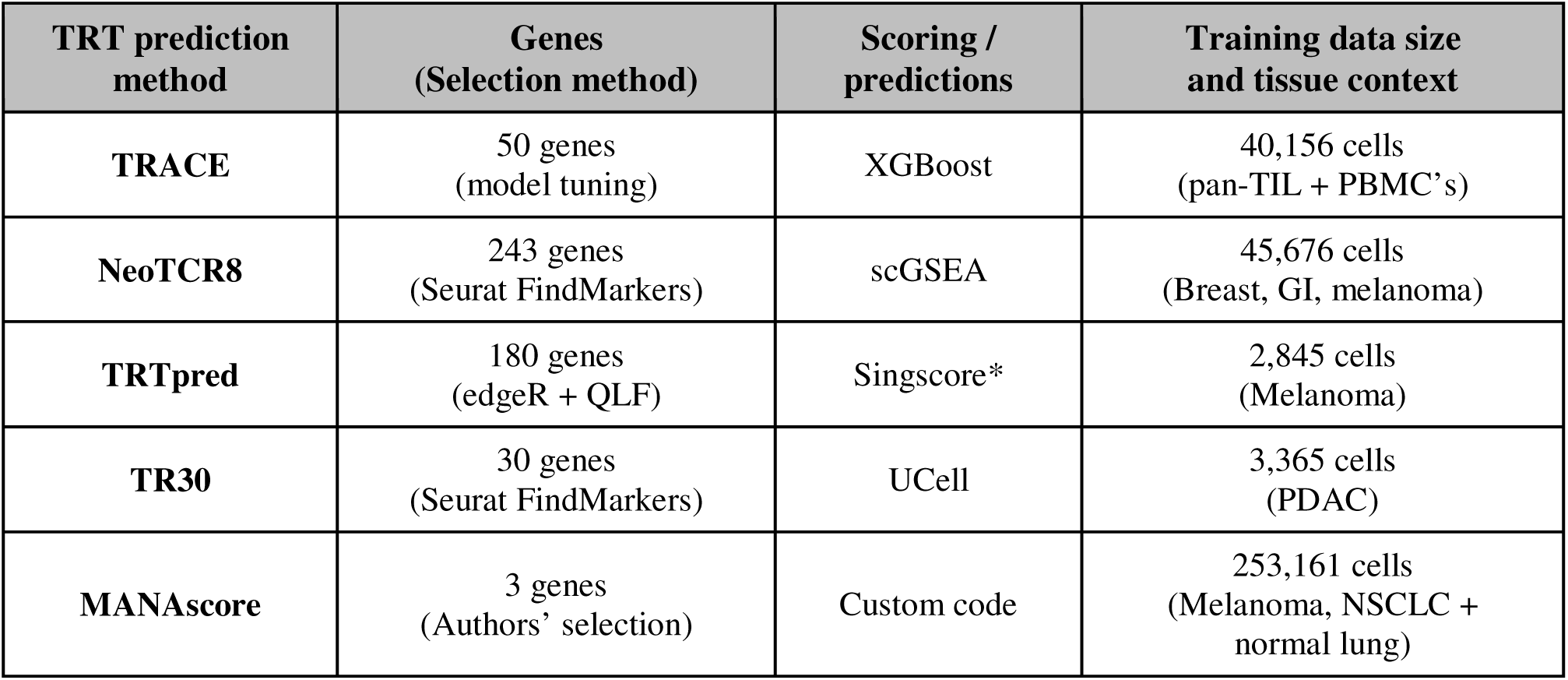
Table with details of other TRT prediction methods implemented and benchmarked against TRACE. *For TRTpred, the final scoring method was not identified, so Singscore was used.

### TRACE precisely identifies antigen-experienced, exhausted CD8^+^ T cells in tumors across multiple indications

We used TRACE to evaluate tumor reactive CD8^+^ cells in hundreds of patient samples across multiple indications - including those containing scRNA but no matched scTCR-sequencing data - and interpreted our findings alongside any available tumor or clinical metadata (Figure 4A). First, we applied TRACE to individual cells from samples in the Zheng *et al.* pan-cancer TIL atlas sequenced using the 10x Genomics platform, consisting of 134 patients across 17 cancer types (31). Cells across all patients were pooled together and grouped by CD8^+^ subtype according to the authors’ annotations. As expected, TRACE scores were highest in the terminal, *GZMK*^+^, and OXPHOS^-^ exhausted T cell (T_ex_) subtypes marked by consistent expression of *HAVCR2*, *CTLA4*, and *CXCL13,* but not in *TCF7^+^* T_ex_, suggesting TRACE successfully identifies cells that have undergone chronic antigen stimulation (Figure 4B).

**Figure 4.**
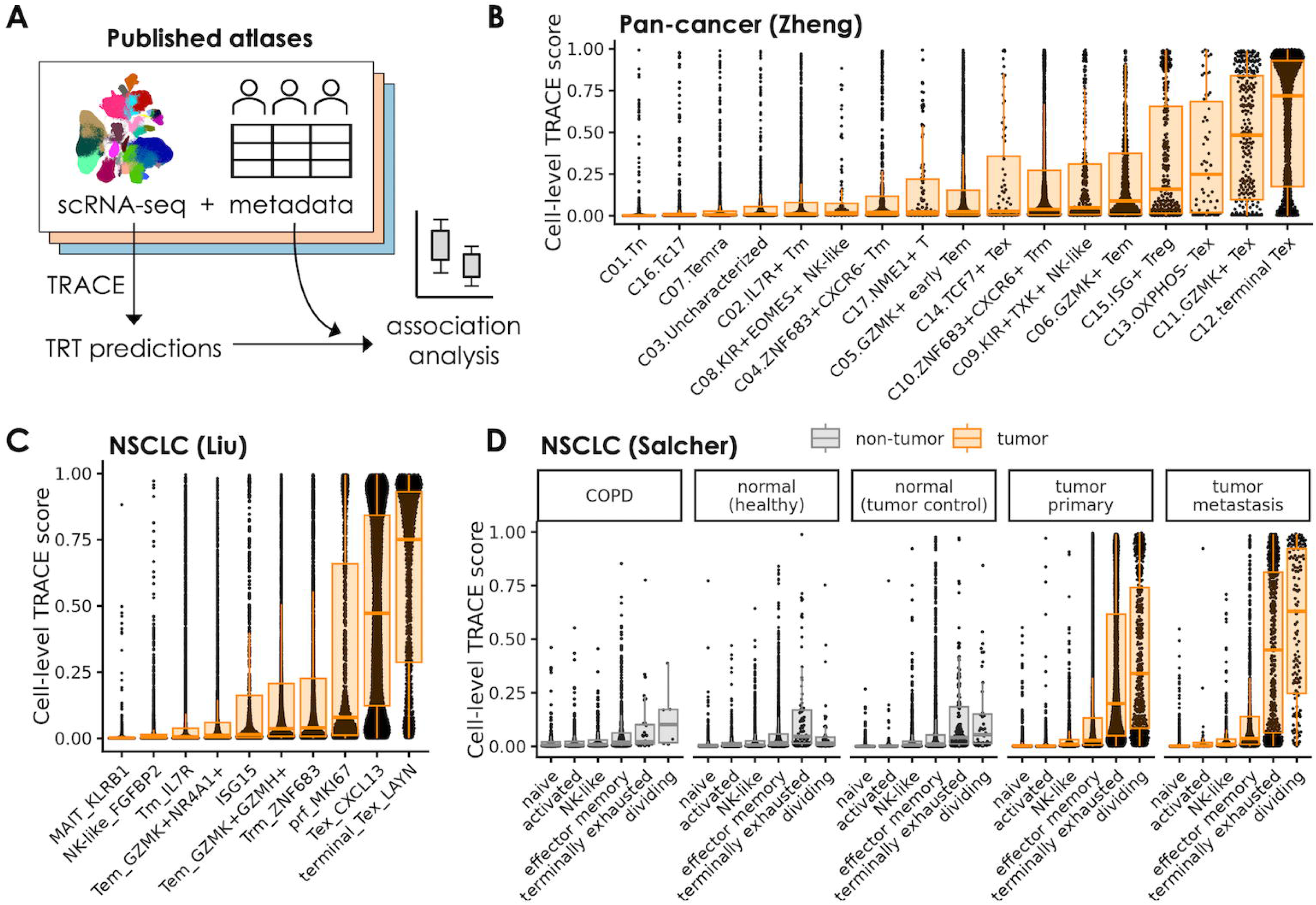
TRACE applied to single cells identifies tumor reactive cells specifically in tumor contexts. **(A)** Application of TRACE to multiple scRNA-seq atlases from the literature. **(B-D)** Distributions of TRACE scores across CD8^+^ cells grouped by CD8^+^ subtype. Tumor cell scores are summarized with orange boxplots, and non-tumor cell scores are summarized with gray boxplots. Cells were subsampled down to 100 cells per sample to ensure relatively even representation of all samples. In B, only CD8^+^ clusters as defined by the original study are included. Scores are shown for the Zheng *et al.* pan-cancer TIL atlas (B), the Liu *et al.* NSCLC atlas (C), and the Salcher *et al.* NSCLC atlas (D).

TRACE produced a similar pattern when applied to the Liu *et al.* NSCLC atlas(37) consisting of 234 NSCLC patients treated with neoadjuvant chemo-immunotherapy, with the two exhausted subtypes (T_ex_ *CXCL13* and terminal T_ex_ *LAYN*) showing the highest scores (Figure 4C). The two exhausted subtypes, along with the proliferative subtype (prf *MKI67*), showed a bimodal distribution with peaks close to either 1 or 0, consistent with confident TRACE predictions of both tumor reactive and exhausted or dividing bystander cells, respectively. The Zheng *et al.* and Liu *et al.* atlases also included some samples with matched scTCR-seq, enabling TRACE scoring at the clone-level. In these datasets, expanded clones (>5 cells) were more likely to score as TRACE^+^ than non- or lowly-expanded clones (Supplementary Figure 4). Given that TRT clones are associated with antigen-mediated expansion at tumor sites, this provides further support that TRACE captures clones that have undergone activation.

We also applied TRACE to the Salcher *et al.* NSCLC atlas(38), which contains samples from 151 NSCLC patients with paired normal samples where available, as well as samples from COPD patients and healthy individuals. Similar to previous datasets, tumor cells in the Salcher *et al.* atlas showed the highest TRACE scores in the terminally exhausted and dividing CD8 subtypes (Figure 4D). This pattern was remarkably consistent across individual datasets and sequencing platforms within the atlas, including 10x Genomics, BD rhapsody, inDrop, Singleron, and Smart-seq 2 (Supplementary Figure 5). Interestingly, non-tumor cells showed a different pattern from tumor cells, with non-tumor cells scoring relatively low for TRACE across all CD8 subtypes in normal control samples as well as healthy and COPD samples. Thus, TRACE uniquely identified cells in exhausted and dividing transcriptomic states within tumor but not in adjacent normal or inflamed non-tumor contexts. TRACE produced a similar result when applied to the Chu *et al.* CRC atlas (39), where we observed higher cell-level TRACE scores in *CXCL13*^+^ CD8^+^ cells from CRC samples compared to cells in a similar subtype within normal controls, healthy individuals, or those with inflammatory bowel disease (IBD) (Supplementary Figure 6). These observations suggest TRACE precisely identifies cells in tumor reactive states and is not merely a measure of general T cell exhaustion. To determine how well TRACE distinguishes tumor-specific reactive states from non-tumor states in comparison to other tumor reactivity prediction tools, we also scored the Salcher *et al.* atlas using the gene sets from TRTpred and NeoTCR8 (24,26). Compared to TRTpred and NeoTCR8, TRACE showed higher separation between exhausted tumor scores and exhausted non-tumor scores, indicating TRACE is highly specific for tumor reactivity relative to other methods (Supplementary Figure 7).

### TRACE identifies tumor subtypes enriched for TRTs

To compare TRACE scores across samples, we defined a sample-level TRACE score (sTRACE score) as the fraction of CD8^+^ T cells in the sample with a TRACE score of 0.5 or higher. In Salcher *et al.*, tumor samples across NSCLC subtypes (LUAD, LUSC, and unspecified) generally exhibited higher sTRACE than non-tumor samples, with false positive sTRACE scores in non-tumor samples of typically < 5% (Figure 5A). sTRACE scores were similar between LUAD and LUSC and were also similar between primary and metastatic tumors within LUAD. To assess if certain driver mutations were associated with higher sTRACE scores in NSCLC, we plotted tumor sTRACE scores from Salcher *et al.* and Liu *et al.* against listed driver mutations. In both studies, *KRAS* mutations were associated with higher sTRACE scores relative to other reported mutations (Figure 5B). High-scoring *KRAS*-mutant samples included both primary and metastatic samples with or without a co-occurring *TP53* mutation.

**Figure 5.**
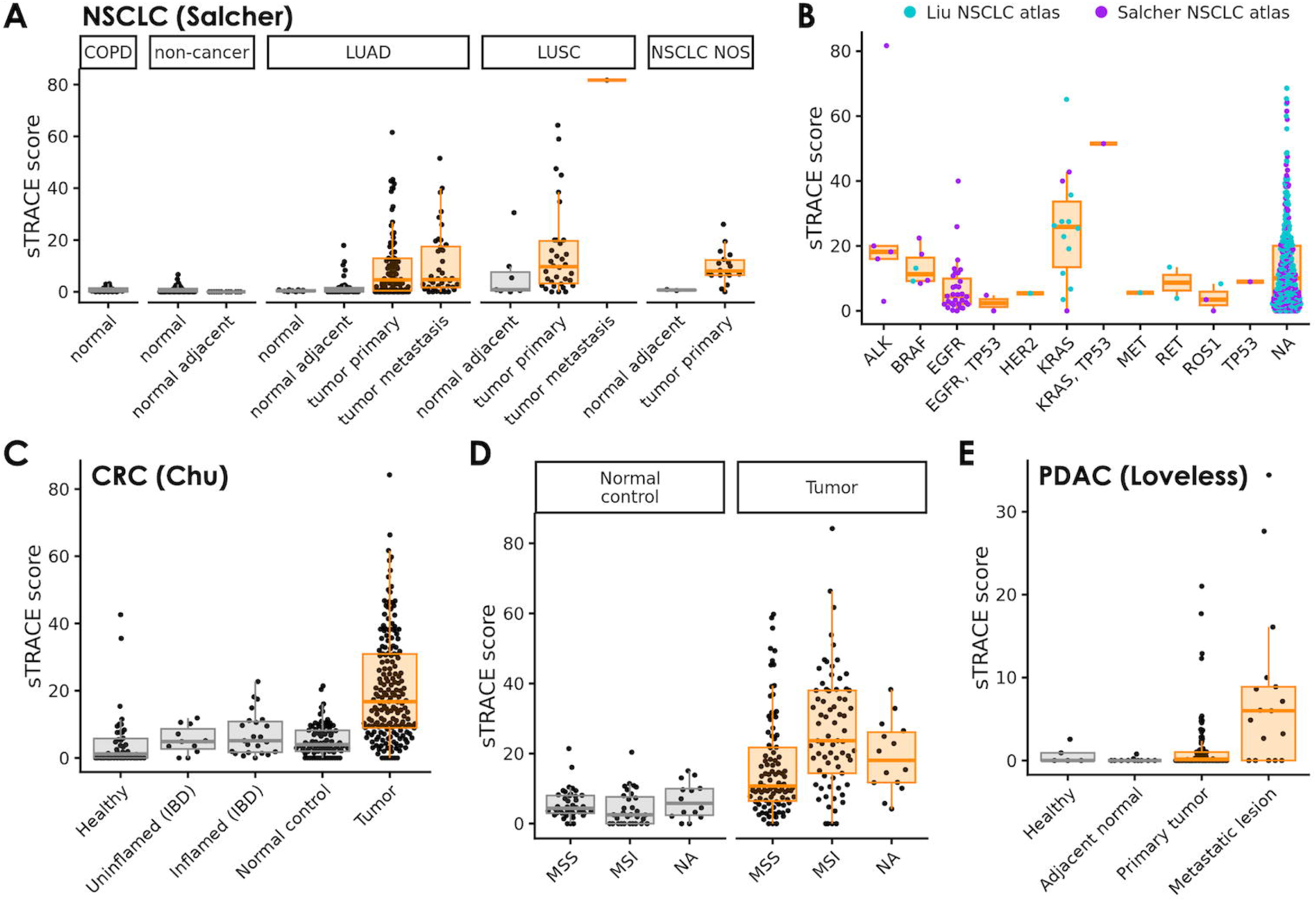
Large-scale survey of tumor reactivity in different cancer types using TRACE. **(A)** sTRACE scores for each NSCLC and non-cancerous sample from the Salcher *et al.* lung atlas. NOS, not otherwise specified. **(B)** sTRACE scores for NSCLC samples grouped by driver gene status. Samples categorized as NA (not available) were not assessed for one or more of the genes listed. **(C)** sTRACE scores for each CRC and non-cancerous sample from the Chu *et al.* CRC atlas. **(D)** sTRACE scores for the CRC and normal control samples from panel C grouped by MSI status. **(E)** sTRACE scores for each PDAC and non-cancerous sample from the Loveless *et al.* PDAC atlas. For all panels, tumor sample scores are summarized with orange boxplots, and non-tumor sample percentages are summarized with gray boxplots.

We similarly applied TRACE to the Chu *et al.* CRC atlas(39) and found the highest sTRACE scores in tumor samples, with lower scores in normal control and healthy samples and in samples from patients with IBD (Figure 5C). Within CRC tumor samples, we further investigated sTRACE scores as a function of MSI status. Colorectal tumors with microsatellite instability (MSI) exhibit higher levels of CD8^+^ T cell infiltration than those that are microsatellite stable (MSS) (40), but the tumor reactivity of these TIL is not well-established. Using TRACE, we found higher sTRACE scores in MSI tumors (p = 5.8×10^-6^; Figure 5D), but no differences between normal control samples from MSS or MSI tumors (p = 0.071). Thus, MSI tumors are immunologically favorable in terms of both CD8^+^ T cell infiltration and tumor reactivity. Finally, we scored CD8^+^ cells in the PDAC atlas compiled by Loveless *et al* (41). In line with the low immunogenicity of PDAC tumors, we observed lower sTRACE scores overall in these samples. Interestingly, the scores were higher in PDAC metastatic lesions (mostly liver) than primary tumors, despite a decrease in CD8^+^ T cell infiltration in these samples.

## 4 Discussion

Tumor reactive CD8^+^ T cells represent a biologically and clinically meaningful subset of TIL, and their frequency and functionality is a key driver of clinical efficacy of immune checkpoint inhibitors (CPI) and adoptive cell therapies (ACT) in immunologically hot tumors. Identifying and estimating the frequency of TRTs in TIL is of interest as a predictive biomarker of response to CPI or ACT regardless of tissue histology. Further, TRTs can be preferentially isolated for *ex vivo* TIL therapy and for discovery of novel TCR sequences specific to tumor associated antigens.

A growing body of research has shown that TRTs occupy distinct transcriptional states enriched for markers of chronic antigen stimulation, tissue residency, and exhaustion like programs (e.g., *CXCL13*, *ENTPD1*/CD39, *PDCD1*, *LAG3*, *TOX*). Prior methods - including gene signature scoring methods (e.g., NeoTCR8, MANAscore), marker based heuristics (e.g., CD39/CD103 co expression), and machine learning classifiers trained on individual studies - have advanced the ability to infer tumor reactivity but are often constrained by their dependence on single datasets, lack of model checkpoint availability, or sensitivity to normalization and platform specific effects. In this study, we introduce TRACE, a clonotype aware, machine learning framework designed to infer tumor reactivity from single cell transcriptomic data while explicitly accounting for technical variability and intra clonal heterogeneity.

Using autologous co culture assays and scRNA/scTCR-seq, we confirm that TRACE predictions correspond closely with experimentally reactive clones, providing a strong functional grounding for model performance. TRACE accurately identifies exhausted or chronically stimulated CD8 T cell populations across NSCLC, CRC, PDAC, and pan cancer datasets, and distinguishes tumor specific activation from bystander exhaustion in non tumor tissues. Associations with clonal expansion and known clinical features (e.g., KRAS mutations, MSI status) underscore biological validity.

By integrating validated TRT and non TRT clonotypes across multiple TIL and PBMC datasets, TRACE addresses key limitations of previous computational approaches and offers a generalizable strategy for identifying tumor reactive T cells in both tissue and blood contexts. Since TRACE is trained on a relatively small set of validated TRT clones even after aggregation of multiple datasets, there is potential for this model to be refined on new datasets such as rare cancer subtypes, or on tumors with atypical TIL transcriptional states following multiple rounds of treatment. Openly sharing the code, model weights, and feature set allows anyone in the community to develop model applications on public atlases or niche datasets. Further, there is potential for this model to be refined with additional autologous co-culture experiments from additional donors across multiple indications, with emphasis on identifying tumor reactive clonotypes with atypical profiles. In conclusion, TRACE is a framework that will be of great interest to the adoptive cell therapy field.

## 5 Materials and Methods

### Data aggregation and cleaning

For all datasets used in this study except two (34,35), we downloaded raw sequencing reads (FASTQ format) to partially mitigate batch effects by ensuring consistent alignment and processing of the reads. When starting with FASTQ files, 10x Genomics Cell Ranger v8.0.1 was used for gene expression quantification, TCR sequence assembly, and extraction of cell barcodes and UMIs. RNA was aligned to a modified version of the GRCh38 reference containing only protein-coding genes, and TCR sequences were aligned to the vdj_GRCh38_alts_ensembl reference. For all datasets, Seurat (version 5.1.0) (42) was used to process feature-barcode matrices, align gene expression and TCR datasets, filter cells, identify and annotate clusters, and perform differential gene expression analyses. Briefly, cells containing fewer than 200 unique genes and cells where the fraction of counts belonging to mitochondrial genes (%mito) exceeded 25% were removed. In addition, cells were scored for how well they expressed S- and G2M-phase gene sets using CellCycleScoring (Seurat). For each cell, a value CCdifference was calculated as the difference between that cell’s S-phase and G2M-phase scores. scRepertoire (43) was used to ensure a maximum of one TCRα and one TCRβ chain per cell.

Where possible, we used experimentally verified TRT and non-TRT labels identified using paired TCRαβ CDR3 sequences provided by the authors. When only TRT TCRβ CDR3 sequences were provided (22), tumor reactivity labels were assigned based on CDR3β sequences alone. For some datasets, we labeled additional clones as likely non-tumor reactive if they contained a CDR3β chain with an annotated target in VDJdb.

After removing clones with unknown TRT status, we further filtered out 1) putative CD4^+^ T cells and 2) samples for which the number of annotated clones was low. CD4^+^ cells were identified based on the relative expressions of *CD4* and *CD8A* after clustering. To cluster the cells, we followed the scTransform workflow (Seurat), regressing %mito and CCdifference. TCR variable and joining genes were removed from the list of genes used for principal component analysis and clustering to avoid clonotype-specific clusters. Dimensionality reduction was performed with 20 principal components, and clustering was performed with resolutions typically between 0.5 and 1.0.

### Data filtering

The assembled dataset contains expression data from 30,560 genes and 40,156 cells representing 16,465 distinct CD8^+^ clones. Prior to model training, the following gene filtering steps were applied to the dataset – 1) selection of genes that are expressed (> 0 count) in at least 10% of cells across the dataset, 2) selection of top 5,000 most highly variable genes using FindVariableGenes (Seurat) and 3) exclusion of mitochondrial, ncRNA, ribosomal and other highly expressed genes (Supplementary methods). 966 genes meeting these criteria were retained for model training. No cells were excluded and the clone labels (TRT vs non-TRT) were not used at any point during filtering.

### Clone-level feature construction

Single-cell expression matrices were summarized to clone-level feature vectors by aggregating expression across all cells assigned to the same clone. Clone expression was summarized using a percentile-based aggregation (75^th^ percentile) to provide a robust clone-level representation while reducing sensitivity to outlier cells. Genes used for model development were drawn from a predefined feature universe defined in the data filtering step. The parameter (75^th^ percentile) for clone collapsing was selected after assessing the results of optimization experiment 1 as described in supplementary methods, where several percentiles were evaluated. In addition, raw counts for genes in each cell were transformed either by expression binning (eb) or log normalization (ln). Expression binning involves binning genes by their non-zero expression count in each cell into *n* equally sized bins (20). All genes with zero counts were grouped into the 0th bin, and each gene’s expression value was represented by their bin count from 0 to n. Expression binning elegantly deals with variation in counts data due to variability in sample batching, library preparations, read depth, or sequencing platforms which frequently plagues single-cell RNAs data analysis. Log normalization involves a transformation of gene expression values within each cell using 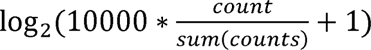 Importantly, neither method requires cell or data set normalization or transformation, such as scTransform. Expression binning was chosen as the default method for pre-processing after clone-level summarization, as it was superior to log normalization (optimization experiment 1) across a range of hyperparameters and resulted in > 5X faster training compared to log normalization.

### Model class and training objective

We used an XGBoost gradient-boosted decision tree classifier. XGBoost was chosen as the top performing model in a head-to-head comparison with randomForest and AdaBoost (Supplementary methods). Because tumor reactive clones represent the clinically relevant positive class and were present at significantly lower prevalence than negative class in the dataset, model selection emphasized positive-class performance, and evaluation focused on positive-class–aware metrics (including PR-AUC and F1 for the positive class), in addition to overall accuracy.

### Cross-validation design and prevention of leakage

To obtain an unbiased estimate of generalization and prevent information leakage from feature selection and hyperparameter tuning, we used nested cross-validation at the clone level. In each outer fold, 20% of the entire dataset was held out for model validation and was not used for model training. 80% of the dataset was used as training data for inner cross-validation. 5-fold cross-validation was performed on the inner fold training data for hyperparameter optimization and feature selection, and the optimal model was re-trained on the full 80% training data and tested on the 20% hold-out data for reporting model performance. The outer fold (training-test-validation) was repeated between 3-50 times for different optimization experiments. During each inner fold cross-validation, TRT positive class and TRT negative class clones were balanced at 10:90 ratio by subsampling negative class clones.

### Hyperparameter optimization

Hyperparameters for XGBoost, randomForest and AdaBoost were optimized within the inner loop using automated search. The TRACE framework supports both randomized search and Bayesian optimization (Optuna); hyperparameter optimization was performed and fold-specific optimal configurations were consolidated using a stability-based selection strategy to reduce overfitting to any single split. The “Optuna” method is default as it had superior performance compared to “random grid search” (Supplementary methods) although both options are available when running TRACE.

### Feature selection and “freezing” of the signature

To improve interpretability and reduce overfitting, we performed cross-validated feature selection using model-derived feature importance computed within the training folds. Importance values were converted to per-fold ranks and aggregated across folds, and a fixed set of genes was selected and then frozen for final model fitting on inner training data and downstream evaluation on hold out validation data. The number of genes to be selected was chosen to be 50 (Supplementary methods). These selected features, in addition to the additional 450 features used to establish the bins - are stored with the trained model and used for consistent feature alignment during inference on new datasets.

### Performance reporting

Model performance was summarized across outer folds (mean ± standard deviation, unless otherwise indicated) and reported on the held-out validation set. Metrics included threshold-independent discrimination (ROC-AUC, PR-AUC) and threshold-dependent classification performance, with particular emphasis on positive-class precision/recall tradeoffs relevant to tumor reactive clone identification.

### Benchmarking

We implemented other published TRT methods from datasets used in this study (NeoTCR8, TRTpred, TR30, and MANAscore) and applied them to our extended dataset.

For the NeoTCR8, TRTpred, and TR30 methods, we 1) scored individual cells in our dataset using their published gene lists and chosen scoring method, 2) calculated clone-level scores from these cell-level scores, 3) trained a logistic regression model using training set clones in each outer fold, and 4) applied these models to predict TRT labels for held-out clones, using the threshold that best separated TRT from non-TRT clones within the training set.

For NeoTCR8, individual cell scores were found using single sample GSEA. To apply NeoTCR8 to multiple training sets without clustering, the clone-wise score was set to be the maximum score found within that clone. Within each seed, the predicted clone labels were found by comparing test set scores to the optimal threshold learned from the training set, maximizing Youden’s J.

For TRTpred, individual cell scores were found using Singscore, as no single method was clearly described. The clone-wise TRTpred score was set to be the maximum score found within that clone. Within each seed, scores were standardized based on the distribution of scores in the training set (mean ± SD), and the same transformation was applied to scores from the test set. The predicted clone labels were identified by comparing the standardized test set scores to the optimal threshold learned from the training set, maximizing accuracy.

For TR30, individual cell scores were found using the UCell package. The clone-wise TR30 score was set to be the mean score found within that clone. Within each seed, the predicted clone labels were identified by comparing test set scores to the optimal threshold learned from the training set, maximizing Youden’s J.

For MANAscore, we used code available in their Github repository to find imputed and non-imputed MANAscores for each cell and called cells scoring higher than the level of the final trough for each score as MANAscore_hi_ cells. Putative tumor reactive clones were defined as those with at least 5 MANAscore_hi_ cells. For a few samples, the provided code was unable to identify the final imputed MANAscore trough; for these samples, we set these thresholds equal to the value of the matched non-imputed MANAscore trough. To compare clone-level TRT scores between methods, we set the clone-level score equal to the maximum imputed MANAscore found within a clone.

### TRACE scoring of public scRNA-seq atlases

Counts were downloaded for each dataset and converted to h5ad where necessary. CD8^+^ cells were selected using the cell type annotations provided by the study authors except in the case of the Loveless *et al.* PDAC atlas, for which CD8^+^ labels were unavailable and were therefore assigned by identifying clusters high for *CD8A/CD8B* expression. TRACE was run in single-cell mode using the config_preprocess_sc.yaml settings with the gene_selection_method set to “all”. For the Zheng *et al*. pan-cancer and Liu *et al.* NSCLC atlases, TRACE was additionally run at the clone level after integrating the scTCR-seq data. TRACE was applied to the raw counts for each atlas, except for the Chu *et al.* CRC atlas, which only contained normalized counts. Resulting TRACE scores were then joined with the patient metadata available from each study. For sample-level sTRACE scores, only samples with ≥ 10 CD8^+^ cells were considered. The Becker *et al*. dataset was removed from the Chu *et al.* CRC atlas since sequencing was performed on single nuclei rather than single cells (44). Additional details on any pre-processing of the atlases can be found in the Supplementary Methods, and the code for score generation is available at https://github.com/ksqtx/trace.

### TIL/tumor autologous co-culture assay

Melanoma tumor fragments were digested and used to develop a melanoma tumor cell line. TIL from tumor fragments were expanded *ex vivo* following protocols described elsewhere and cryopreserved (45,46). After thaw, expanded TIL were rested for 24 h prior to co-culture with the matched autologous melanoma cell line at an E:T ratio of 0.5:1 (300,000 T cells and 600,000 tumor cells per well) for 20 h. To evaluate MHC-mediated T cell activation, an MHC-block control condition was used, which involved incubating tumor cells with anti-MHC-I (168 ug/mL, Ultra-LEAF clone W6/32, BioLegend) and anti-MHC-II (50 ug/mL, BD Pharmingen clone Tu39, BD) at 37 °C for 1 h prior to co-culture.

Prior to sequencing, T cells from tumor starting material and from autologous co-cultures underwent dead cell removal (#130-090-101, Miltenyi Biotec) followed by T cell enrichment using CD3 microbeads (EasySep Human CD3 #17851, STEMCELL Technologies). Sequencing was performed using the 10x Genomics platform and RNA and VDJ libraries were prepared by the Chromium Next GEM Single Cell 5’v2 HT Reagent kit according to the manufacturer’s protocol (10x Genomics).

Expt. TIL clones were identified as any clone with at least 1 cell and 5% of cells in a *TNFRSF9*-high transcriptional state in any replicate.

## 7 Author Contributions

DM: Conceptualization, Data curation, Methodology, Software, Visualization, Writing – original draft, Writing – review & editing. JD: Data curation, Methodology, Software, Writing – review & editing. AED: Data curation, Methodology, Visualization, Writing – original draft, Writing – review & editing. AY: Conceptualization, Methodology, Writing – review & editing. MG: Investigation, Writing – review & editing. MT: Investigation, Writing – review & editing. GJM: Investigation, Supervision, Writing – review & editing. SC: Investigation, Writing – review & editing. CC: Investigation, Writing – review & editing. CHY: Methodology, Investigation, Writing – review & editing. KW: Supervision, Writing – review & editing. MJB: Supervision, Writing – review & editing. DS: Conceptualization, Data curation, Methodology, Project administration, Software, Supervision, Writing – original draft, Writing – review & editing.

## 8 Acknowledgements

We gratefully acknowledge the patients and donors whose tumor specimens made this work possible. We would like to acknowledge Shreya Patil for her help in data processing.

## 10 Data Availability Statement

Details about the training and atlas datasets can be found in the supplementary materials file.

## 11 Code availability

The TRACE model, model checkpoint weights, and instructions for its use can be found at https://github.com/ksqtx/trace.

## References

1. Tumeh PC, Harview CL, Yearley JH, Shintaku IP, Taylor EJM, Robert L, Chmielowski B, Spasic M, Henry G, Ciobanu V, et al. PD-1 blockade induces responses by inhibiting adaptive immune resistance. Nature (2014) 515:568–571. doi: 10.1038/nature13954

2. Rosenberg SA, Restifo NP, Yang JC, Morgan RA, Dudley ME. Adoptive cell transfer: a clinical path to effective cancer immunotherapy. Nature Reviews Cancer (2008) 8:299–308. doi: 10.1038/nrc2355

3. Sade-Feldman M, Yizhak K, Bjorgaard SL, Ray JP, Boer CG de, Jenkins RW, Lieb DJ, Chen JH, Frederick DT, Barzily-Rokni M, et al. Defining T Cell States Associated with Response to Checkpoint Immunotherapy in Melanoma. Cell (2018) 175:998–1013.e20. doi: 10.1016/j.cell.2018.10.038

4. Litchfield K, Reading JL, Puttick C, Thakkar K, Abbosh C, Bentham R, Watkins TBK, Rosenthal R, Biswas D, Rowan A, et al. Meta-analysis of tumor- and T cell-intrinsic mechanisms of sensitization to checkpoint inhibition. Cell (2021) 184:596–614. doi: 10.1016/j.cell.2021.01.002

5. Bruni D, Angell HK, Galon J. The immune contexture and Immunoscore in cancer prognosis and therapeutic efficacy. Nature Reviews Cancer (2020) 20:662–680. doi: 10.1038/s41568-020-0285-7

6. Simoni Y, Becht E, Fehlings M, Loh CY, Koo S-L, Teng KWW, Yeong JPS, Nahar R, Zhang T, Kared H, et al. Bystander CD8+ T cells are abundant and phenotypically distinct in human tumour infiltrates. Nature (2018) 557:575–579. doi: 10.1038/s41586-018-0130-2

7. Meier SL, Satpathy AT, Wells DK. Bystander T cells in cancer immunology and therapy. Nature Cancer (2022) 3:143–155. doi: 10.1038/s43018-022-00335-8

8. Duhen T, Duhen R, Montler R, Moses J, Moudgil T, Miranda NF de, Goodall CP, Blair TC, Fox BA, McDermott JE, et al. Co-expression of CD39 and CD103 identifies tumor-reactive CD8 T cells in human solid tumors. Nature Communications (2018) 9:2724. doi: 10.1038/s41467-018-05072-0

9. Thommen DS, Koelzer VH, Herzig P, Roller A, Trefny M, Dimeloe S, Kiialainen A, Hanhart J, Schill C, Hess C, et al. A transcriptionally and functionally distinct PD-1+ CD8+ T cell pool with predictive potential in non-small-cell lung cancer treated with PD-1 blockade. Nature Medicine (2018) 24:994–1004. doi: 10.1038/s41591-018-0057-z

10. Chow A, Uddin FZ, Liu M, Dobrin A, Nabet BY, Mangarin L, Lavin Y, Rizvi H, Tischfield SE, Quintanal-Villalonga A, et al. The ectonucleotidase CD39 identifies tumor-reactive CD8+ T cells predictive of immune checkpoint blockade efficacy in human lung cancer. Immunity (2023) 56:93–106.e6. doi: 10.1016/j.immuni.2022.12.001

11. Yost KE, Satpathy AT, Wells DK, Qi Y, Wang C, Kageyama R, McNamara KL, Granja JM, Sarin KY, Brown RA, et al. Clonal replacement of tumor-specific T cells following PD-1 blockade. Nature Medicine (2019) 25:1251–1259. doi: 10.1038/s41591-019-0522-3

12. Barras D, Ghisoni E, Chiffelle J, Orcurto A, Dagher J, Fahr N, Benedetti F, Crespo I, Grimm AJ, Morotti M, et al. Response to tumor-infiltrating lymphocyte adoptive therapy is associated with preexisting CD8 ^+^ T-myeloid cell networks in melanoma. Science Immunology (2024) 9: doi: 10.1126/sciimmunol.adg7995

13. Scheper W, Kelderman S, Fanchi LF, Linnemann C, Bendle G, Rooij MAJ de, Hirt C, Mezzadra R, Slagter M, Dijkstra K, et al. Low and variable tumor reactivity of the intratumoral TCR repertoire in human cancers. Nature Medicine (2019) 25:89–94. doi: 10.1038/s41591-018-0266-5

14. Miller BC, Sen DR, Abosy RA, Bi K, Virkud YV, LaFleur MW, Yates KB, Lako A, Felt K, Naik GS, et al. Subsets of exhausted CD8+ T cells differentially mediate tumor control and respond to checkpoint blockade. Nature Immunology (2019) 20:326–336. doi: 10.1038/s41590-019-0312-6

15. Liu B, Zhang Y, Wang D, Hu X, Zhang Z. Single-cell meta-analyses reveal responses of tumor-reactive CXCL13+ T cells to immune-checkpoint blockade. Nature Cancer (2022) 3:1123–1136. doi: 10.1038/s43018-022-00433-7

16. Krishna S, Lowery FJ, Copeland AR, Bahadiroglu E, Mukherjee R, Jia L, Anibal JT, Sachs A, Adebola SO, Gurusamy D, et al. Stem-like CD8 T cells mediate response of adoptive cell immunotherapy against human cancer. Science (2020) 370:1328–1334. doi: 10.1126/science.abb9847

17. Lowery FJ, Goff SL, Gasmi B, Parkhurst MR, Ratnam NM, Halas HK, Shelton TE, Langhan MM, Bhasin A, Dinerman AJ, et al. Neoantigen-specific tumor-infiltrating lymphocytes in gastrointestinal cancers: a phase 2 trial. Nature Medicine (2025) 31:1994–2003. doi: 10.1038/s41591-025-03627-5

18. Zacharakis N, Chinnasamy H, Black M, Xu H, Lu Y-C, Zheng Z, Pasetto A, Langhan M, Shelton T, Prickett T, et al. Immune recognition of somatic mutations leading to complete durable regression in metastatic breast cancer. Nature Medicine (2018) 24:724–730. doi: 10.1038/s41591-018-0040-8

19. Berg JH van den, Heemskerk B, Rooij N van, Gomez-Eerland R, Michels S, Zon M van, Boer R de, Bakker NAM, Jorritsma-Smit A, Buuren MM van, et al. Tumor infiltrating lymphocytes (TIL) therapy in metastatic melanoma: boosting of neoantigen-specific T cell reactivity and long-term follow-up. Journal for ImmunoTherapy of Cancer (2020) 8:e000848. doi: 10.1136/jitc-2020-000848

20. Seliktar-Ofir S, Merhavi-Shoham E, Itzhaki O, Yunger S, Markel G, Schachter J, Besser MJ. Selection of Shared and Neoantigen-Reactive T Cells for Adoptive Cell Therapy Based on CD137 Separation. Frontiers in Immunology (2017) 8: doi: 10.3389/fimmu.2017.01211

21. Ye Q, Song D-G, Poussin M, Yamamoto T, Best A, Li C, Coukos G, Powell DJ. CD137 Accurately Identifies and Enriches for Naturally Occurring Tumor-Reactive T Cells in Tumor. Clinical Cancer Research (2014) 20:44–55. doi: 10.1158/1078-0432.ccr-13-0945

22. Caushi JX, Zhang J, Ji Z, Vaghasia A, Zhang B, Hsiue EHC, Mog BJ, Hou W, Justesen S, Blosser R, et al. Transcriptional programs of neoantigen-specific TIL in anti-PD-1-treated lung cancers. Nature (2021) 596:126–132. doi: 10.1038/s41586-021-03752-4

23. Hanada K ichi, Zhao C, Gil-Hoyos R, Gartner JJ, Chow-Parmer C, Lowery FJ, Krishna S, Prickett TD, Kivitz S, Parkhurst MR, et al. A phenotypic signature that identifies neoantigen-reactive T cells in fresh human lung cancers. Cancer Cell (2022) 40:479–493.e6. doi: 10.1016/j.ccell.2022.03.012

24. Lowery FJ, Krishna S, Yossef R, Parikh NB, Chatani PD, Zacharakis N, Parkhurst MR, Levin N, Sindiri S, Sachs A, et al. Molecular signatures of antitumor neoantigen-reactive T cells from metastatic human cancers. Science (2022) 375:877–884. doi: 10.1126/science.abl5447

25. Oliveira G, Stromhaug K, Klaeger S, Kula T, Frederick DT, Le PM, Forman J, Huang T, Li S, Zhang W, et al. Phenotype, specificity and avidity of antitumour CD8+ T cells in melanoma. Nature (2021) 596:119–125. doi: 10.1038/s41586-021-03704-y

26. Pétremand R, Chiffelle J, Bobisse S, Perez MAS, Schmidt J, Arnaud M, Barras D, Lozano-Rabella M, Genolet R, Sauvage C, et al. Identification of clinically relevant T cell receptors for personalized T cell therapy using combinatorial algorithms. Nature Biotechnology (2025) 43:323–328. doi: 10.1038/s41587-024-02232-0

27. Zeng Z, Zhang T, Zhang J, Li S, Connor S, Zhang B, Zhao Y, Wilson J, Singh D, Kulikauskas R, et al. A minimal gene set characterizes TIL specific for diverse tumor antigens across different cancer types. Nature Communications (2025) 16: doi: 10.1038/s41467-024-55059-3

28. Tan CL, Lindner K, Boschert T, Meng Z, Ehrenfried AR, Roia AD, Haltenhof G, Faenza A, Imperatore F, Bunse L, et al. Prediction of tumor-reactive T cell receptors from scRNA-seq data for personalized T cell therapy. Nature Biotechnology (2025) 43:134–142. doi: 10.1038/s41587-024-02161-y

29. Li H, Leun AM van der, Yofe I, Lubling Y, Gelbard-Solodkin D, Akkooi ACJ van, Braber M van den, Rozeman EA, Haanen JBAG, Blank CU, et al. Dysfunctional CD8 T Cells Form a Proliferative, Dynamically Regulated Compartment within Human Melanoma. Cell (2020) 181:747. doi: 10.1016/j.cell.2020.04.017

30. Krishna C, DiNatale RG, Kuo F, Srivastava RM, Vuong L, Chowell D, Gupta S, Vanderbilt C, Purohit TA, Liu M, et al. Single-cell sequencing links multiregional immune landscapes and tissue-resident T cells in ccRCC to tumor topology and therapy efficacy. Cancer Cell (2021) 39:662–677.e6. doi: 10.1016/j.ccell.2021.03.007

31. Zheng L, Qin S, Si W, Wang A, Xing B, Gao R, Ren X, Wang L, Wu X, Zhang J, et al. Pan-cancer single-cell landscape of tumor-infiltrating T cells. Science (2021) 374: doi: 10.1126/science.abe6474

32. Meng Z, Ehrenfried AR, Tan CL, Steffens LK, Kehm H, Zens S, Lauenstein C, Paul A, Schwab M, Förster JD, et al. Transcriptome-based identification of tumor-reactive and bystander CD8 ^+^ T cell receptor clonotypes in human pancreatic cancer. Science Translational Medicine (2023) 15: doi: 10.1126/scitranslmed.adh9562

33. Gao S, Wu Z, Arnold B, Diamond C, Batchu S, Giudice V, Alemu L, Raffo DQ, Feng X, Kajigaya S, et al. Single-cell RNA sequencing coupled to TCR profiling of large granular lymphocyte leukemia T cells. Nat Commun (2022) 13:1982. doi: 10.1038/s41467-022-29175-x

34. Genomics 10x. Human PBMC from a Healthy Donor, 10k cells (v2). (2020) https://www.10xgenomics.com/datasets/human-pbmc-from-a-healthy-donor-10-k-cells-v-2-2-standard-5-0-0

35. Ogura H, Gohda J, Lu X, Yamamoto M, Takesue Y, Son A, Doi S, Matsushita K, Isobe F, Fukuda Y, et al. Dysfunctional Sars-CoV-2-M protein-specific cytotoxic T lymphocytes in patients recovering from severe COVID-19. Nat Commun (2022) 13:7063. doi: 10.1038/s41467-022-34655-1

36. Goncharov M, Bagaev D, Shcherbinin D, Zvyagin I, Bolotin D, Thomas PG, Minervina AA, Pogorelyy MV, Ladell K, McLaren JE, et al. VDJdb in the pandemic era: a compendium of T cell receptors specific for SARS-CoV-2. Nat Methods (2022) 19:1017–1019. doi: 10.1038/s41592-022-01578-0

37. Liu Z, Yang Z, Wu J, Zhang W, Sun Y, Zhang C, Bai G, Yang L, Fan H, Chen Y, et al. A single-cell atlas reveals immune heterogeneity in anti-PD-1-treated non-small cell lung cancer. Cell (2025) 188:3081–3096.e19. doi: 10.1016/j.cell.2025.03.018

38. Salcher S, Sturm G, Horvath L, Untergasser G, Kuempers C, Fotakis G, Panizzolo E, Martowicz A, Trebo M, Pall G, et al. High-resolution single-cell atlas reveals diversity and plasticity of tissue-resident neutrophils in non-small cell lung cancer. Cancer Cell (2022) 40:1503–1520.e8. doi: 10.1016/j.ccell.2022.10.008

39. Chu X, Li X, Zhang Y, Dang G, Miao Y, Xu W, Wang J, Zhang Z, Cheng S. Integrative single-cell analysis of human colorectal cancer reveals patient stratification with distinct immune evasion mechanisms. Nat Cancer (2024) 5:1409–1426. doi: 10.1038/s43018-024-00807-z

40. Wang Q, Yu M, Zhang S. The characteristics of the tumor immune microenvironment in colorectal cancer with different MSI status and current therapeutic strategies. Front Immunol (2025) 15:1440830. doi: 10.3389/fimmu.2024.1440830

41. Loveless IM, Kemp SB, Hartway KM, Mitchell JT, Wu Y, Zwernik SD, Salas-Escabillas DJ, Brender S, George M, Makinwa Y, et al. Human Pancreatic Cancer Single-Cell Atlas Reveals Association of CXCL10+ Fibroblasts and Basal Subtype Tumor Cells. Clin Cancer Res (2025) 31:756–772. doi: 10.1158/1078-0432.ccr-24-2183

42. Hao Y, Stuart T, Kowalski MH, Choudhary S, Hoffman P, Hartman A, Srivastava A, Molla G, Madad S, Fernandez-Granda C, et al. Dictionary learning for integrative, multimodal and scalable single-cell analysis. Nat Biotechnol (2024) 42:293–304. doi: 10.1038/s41587-023-01767-y

43. Yang Q, Safina KR, Nguyen KDQ, Tuong ZK, Borcherding N. scRepertoire 2: Enhanced and efficient toolkit for single-cell immune profiling. PLoS Comput Biol (2025) 21:e1012760. doi: 10.1371/journal.pcbi.1012760

44. Becker WR, Nevins SA, Chen DC, Chiu R, Horning AM, Guha TK, Laquindanum R, Mills M, Chaib H, Ladabaum U, et al. Single-cell analyses define a continuum of cell state and composition changes in the malignant transformation of polyps to colorectal cancer. Nat Genet (2022) 54:985–995. doi: 10.1038/s41588-022-01088-x

45. Schlabach MR, Lin S, Collester ZR, Wrocklage C, Shenker S, Calnan C, Xu T, Gannon HS, Williams LJ, Thompson F, et al. Rational design of a SOCS1-edited tumor-infiltrating lymphocyte therapy using CRISPR/Cas9 screens. J Clin Investig (2023) 133:e163096. doi: 10.1172/jci163096

46. Mercier IL, Monteiro D, Halpin-Veszeleiova K, Wong K, Dodson A, Martinez GJ, Matos D, Hamza B, Yeri A, McKenney S, et al. Dual-inactivation of Regnase-1 and SOCS1 rewires exhausted CD8+ T cell fate to enhance anti-tumor functionality. (2026) doi: 10.64898/2026.01.21.700812

