## Supplementary_material for "Tumor reactivity assessment using clonal expression (TRACE) reveals tumor reactive CD8^+^ T cell heterogeneity across solid tumors"

### 1 Extended methods

**Software implementation and availability**

All analyses were performed using TRACE, a Python (≥3.11) command-line and library pipeline for supervised classification of single-cell RNA-seq data in either clone-summarized or cell-level mode.

**Data pre-processing**

Prior to TRACE analysis, the assembled dataset was filtered for additional genes to exclude mitochondrial, ribosomal, lncRNA and other structural genes using the following regex pattern
"^MT-", "^RPS", "^RPL", "^HSP", "^TUB", "^H[1234]", "MALAT1", "NEAT1", "EEF1A1", "ACTB", "GAPDH", "TMSB4X". This step is only done during training and is not required for applying TRACE on test datasets for prediction.

**Input data formats and metadata extraction**

TRACE ingests AnnData objects (.h5ad) via Scanpy, and can also read 10x Genomics HDF5 (.h5) using scanpy.read_10x_h5() (used for prediction and analysis-mode support). For .h5ad inputs, cell identifiers are taken from adata.obs.index, gene identifiers from adata.var.index, and the expression matrix from the configured AnnData layer (default: X). Clone identifiers and labels are extracted from adata.obs using either user-specified columns (via config) or auto-detection:

- Clone IDs: specified clone_id_column if present; else auto-detects common names (e.g., clone_ID), else (if allowed) falls back to per-cell “clone” IDs.
- Treatment/condition labels: specified label_column if present; else auto-detects common names (e.g., trt_label), else (if allowed) generates default labels.

A clone to treatment mapping is constructed when labels are available and is used downstream so that clone-level examples inherit their treatment label.

**Quality control and filtering**

Quality control is optionally performed prior to gene selection and summarization. The pipeline computes per-cell and per-gene sparsity and count summaries, clone size distributions, and mitochondrial content (genes whose names begin with MT-/MT). Filtering is applied sequentially:

- Cell filtering by detected genes per cell (≥ min_genes_per_cell; optionally ≤ max_genes_per_cell)
- Gene filtering by number of cells expressing the gene (≥ min_cells_per_gene; optionally ≤ max_cells_per_gene)
- Clone filtering by clone size (≥ min_cells_per_clone)
- Mitochondrial filtering by maximum mitochondrial fraction per cell (default 20%)

Composite quality thresholding (optional): if quality_threshold < 1.0, TRACE computes a 0–1 composite cell-quality score (weighted combination of gene-count score, log-expression score, and mitochondrial score) and removes cells below threshold. (For TRACE model training in this manuscript, no quality filtering was done beyond what was described in method section i.e. quality_threshold was set to 1.0)

Default preprocessing parameters are configurable and documented in config/TRACE_default.yaml (e.g., min_genes_per_cell=100, min_cells_per_clone=1, summarization percentile =75).

**Gene selection**

After QC filtering, genes are selected using one of:

1. All genes (no feature filtering)
2. Highly variable genes (HVGs) computed with Scanpy sc.pp.highly_variable_genes(..., flavor="seurat_v3"), selecting the n_top_genes
3. Variance-based selection, selecting the n_top_genes by variance across cells

Selected genes define the feature space passed into subsequent clone summarization and model training.

**Clone summarization (optional) and cell-level mode**

TRACE supports two analysis modes:

1. Clone-summarized mode (default): cells are grouped by clone ID and each clone is represented by a single feature vector, computed per gene as the mean, median, or percentile (default: 75th percentile) of expression across cells within the clone. Clones with fewer than min_cells_per_clone cells are excluded. A heuristic “clone confidence score” is computed from (i) clone size and (ii) mean per-gene variance within the clone, and stored as metadata.
2. Cell-level mode: clone summarization is skipped and each cell is treated as an independent sample. Cell IDs can optionally be taken from a specified cell_id_column in obs metadata.

**Fold-specific preprocessing (training-time transforms)**

During training and prediction, TRACE can apply the same fold preprocessing chosen at training time:

- Log normalization: per-sample normalization to 10,000 total counts followed by log2 transform, i.e. $\log_{2} (10000*\frac{count}{sum\left( counts \right)}+1)$, with values rounded to 4 decimals for performance.
- Expression binning: per-sample equal-frequency binning of non-zero expression values into n_expression_bins quantile bins (zeros remain 0). This is performed independently per sample to avoid leakage across samples.

**Model training: standard cross-validation with held-out test set**

For standard (non-nested) training, TRACE:

1. Constructs X and y from clone (or cell) feature vectors and labels and applies consistent label encoding.
2. Optionally applies class balancing by reproducible subsampling to a user-specified target ratio (e.g., "yes:no_10:90"), after verifying feasibility for the requested CV and split sizes.
3. Splits data into train+validation (80%) and a held-out test set (20%), using stratified splitting when possible.
4. Performs k-fold cross-validation (default: stratified k=5) on the train+validation partition. Within each fold, fold preprocessing is applied to the fold’s train/validation data prior to fitting. Fold performance is tracked and per-fold confusion matrices and threshold-sweep summaries are saved.
5. Retrains the model on the full train+validation data and evaluates once on the held-out test set. Comprehensive metrics (below) are computed on the held-out evaluation.

Supported algorithms include XGBoost, Random Forest, and AdaBoost; for XGBoost, TRACE automatically selects GPU acceleration when available, otherwise falling back to CPU.

**Optional feature selection from cross-validation**

When enabled, TRACE performs CV-based gene selection using model-derived feature importance across folds:

1. Per-fold feature importance is converted to ranks.
2. Ranks are aggregated across folds (median or mean aggregation).
3. The final gene set is the top top_n_genes by aggregated rank.

Importantly, TRACE stores both (i) the full training gene universe used during preprocessing/CV (all_training_genes) and (ii) the final selected gene subset used for model fitting, enabling consistent prediction-time alignment.

**Hyperparameter optimization and nested cross-validation (optional)**

TRACE supports hyperparameter tuning via grid search, random search, or Optuna Bayesian optimization, using stratified cross-validation and a configurable objective function (e.g., accuracy, F1, precision/recall, ROC-AUC). For nested CV, TRACE implements:

- An outer split into inner (80%) and a final untouched validation set (20%).
- An outer StratifiedKFold loop (default outer_cv_folds=10), where for each outer fold:
- Fold preprocessing is applied.
- Hyperparameters are tuned on the outer-fold training portion only (inner CV), then features are ranked and the top top_n_genes are selected.
- The model is trained on the outer-fold training portion and evaluated on the outer-fold test portion.
- Across outer folds, gene ranks are aggregated (rank-based) to define a stable feature set; hyperparameters are selected across folds using a configurable strategy (best score / most stable within tolerance / most frequent), with optional global retuning on the full inner dataset.
- The final model is trained on the full inner dataset using stable features, and evaluated once on the untouched validation set.

**Model evaluation and reporting**

TRACE computes a comprehensive set of performance metrics, including:

- Threshold-independent: ROC-AUC and PR-AUC (binary or weighted one-vs-rest for multiclass)
- Threshold-dependent: accuracy, precision, recall, weighted F1, F_0.5, balanced accuracy, confusion matrix
- Positive-class focused (binary, optional): sensitivity, specificity, Youden’s J, Matthews correlation coefficient, F_2, F_0.5, and PR-AUC for the positive class

TRACE additionally performs systematic threshold sweeps and records best thresholds per metric. Outputs can include fold-level prediction CSVs and JSON summaries suitable for downstream reporting.

**Prediction on new datasets and feature alignment**

For inference, TRACE enforces alignment of incoming datasets to the trained model’s feature space:

1. Input gene vectors are aligned to the model’s stored all_training_genes (missing genes are padded with zeros; extra genes are ignored).
2. The model’s stored preprocessing method (log normalization or expression binning) is applied.
3. If the model was trained with a selected-gene subset, features are further projected onto the selected gene list in the correct order.

For large .h5ad inputs (including remote S3/HTTPS sources), TRACE supports chunked prediction, processing the dataset in fixed-size chunks and writing predictions incrementally to CSV to bound memory use.

**Reproducibility and experiment tracking**

All runs are parameterized through YAML configuration (or CLI options), including random seeds and preprocessing/training choices. TRACE logs structured metadata (e.g., selected genes, preprocessing parameters, and training gene universes) inside model artifacts to ensure reproducible prediction-time preprocessing. Optional MLflow integration enables centralized tracking of runs, metrics, and model artifacts, including checkpointing and model registry support.

**TRACE scoring of public scRNA-seq atlases**

scRNA-seq raw counts, metadata, and TCR annotations from the Zheng *et al.* pan-cancer TIL atlas were downloaded from Zenodo. These data consisted of samples collected by the authors and data the authors obtained from other sources. For samples collected by the authors, Seurat objects containing CD8^+^ T cells and their metadata were available and merged with TCR data. External datasets were provided as SingleCellExperiment objects containing CD8^+^ T cells with corresponding metadata and these were converted to Seurat objects. The datasets were subsequently converted to .h5ad Anndata objects. TRACE was run in single-cell mode on the combined datasets, and additionally in clone-level mode for the datasets the authors generated with TCR information available.

The scRNA-seq raw counts, scRNA-seq metadata, and TCR annotations for the Liu et al. NSCLC atlas were obtained from GEO. Using the Seurat package (v5.3.1) in R (v4.4.2), The scRNA-seq counts and metadata were combined into a Seurat object and were subset for CD8^+^ T cells using the sub_cell_type labels. The AES gene symbol was renamed to TLE5 to match the gene symbol used in TRACE model. Then TCR annotations were added to the metadata, and the Seurat object was converted to h5ad for scoring with TRACE using the SeuratDisk package (v0.0.0.9021). TRACE was run in both single-cell mode (using config_preprocess_sc.yaml) and clone-level mode (config_preprocess_clone.yaml) with the gene_selection_method switched to "all". A small subset of clones were split across chunks, and the clone-level scores for these clones were averaged between the two chunks. The resulting TRACE predictions were annotated with metadata provided by Liu *et al.*

The NSCLC atlas from Salcher *et al.* was retrieved by downloading the build_atlas_results.tar.xz file from Zenodo and locating the file named 20_build_atlas/annotate_datasets/35_final_atlas/artifacts/ full_atlas_annotated.h5ad. The "raw_counts" layer of the h5ad file was then scored with TRACE in single-cell mode (using config_preprocess_sc.yaml) with the gene_selection_method switched to "all". The target-cells option was used to select for CD8^+^ T cells. The resulting TRACE predictions were then annotated with metadata provided by Salcher et al. as well as metadata manually curated as part of this study.

The CRC atlas from Chu *et al.* was retrieved by downloading the adata_AllAnnotated.h5ad file from https://doi.org/10.6084/m9.figshare.25323397. The "X" layer of the h5ad file was then scored with TRACE in single-cell mode (using config_preprocess_sc.yaml) with the gene_selection_method switched to "all". The target-cells option was used to select for CD8 cells. The resulting TRACE predictions were then annotated with metadata provided by Chu et al.

The PDAC atlas from Loveless *et al.* was available as a Seurat object from Zenodo. Since CD8^+^ T cell labels were unavailable, CD8^+^ T cells were selected by first selecting for "TNK" cells using the authors' annotations, re-normalizing and clustering the TNK subset, and then identifying subclusters with relatively high *CD8A*/*CD8B* expression. After subsetting for CD8^+^ T cells, the Seurat object was converted to h5ad for scoring with TRACE. TRACE was run in single-cell mode (using config_preprocess_sc.yaml) with the gene_selection_method switched to "all". The resulting TRACE predictions were then annotated with metadata provided by Loveless et al.

### 2 Optimization Experiments

We conducted two systematic optimization experiments (Supplementary Table 1) to evaluate the impact of different methodological choices on model performance.

All experiments used the same input dataset and employed nested cross-validation with feature selection enabled using median aggregation. Each experiment varied one or more key parameters while holding others constant, as described below.

**Supplementary Table 1: Summary of optimization experiments performed to tune TRACE parameters**

| **Experiment** | **Description** | **Parameter space** |
| --- | --- | --- |
| 1 | Feature Number and Percentile (for clone summarization) Sweep | Feature number: 10, 25, 50, 75, 100, 200 500  Percentile: 25, 50, 75, 100  Preprocessing method: (log transformation, expression binning)  Tuning method: Optuna (Bayesian), Random grid search |
| 2 | ML algorithm comparison | XGBoost, adaBoost, randomForest |

##### Experiment 1: Feature Selection Parameter Sweep

We systematically varied two feature selection parameters: the number of top genes selected (top_n_genes) and the percentile threshold for gene selection. The experiment evaluated 4 percentile values (25, 50, 75, 100) and 7 top_n_genes values (10, 25, 50, 75, 100, 200, 500) across 20 random seeds (seeds 1-20), resulting in 5,600 total training runs (4 × 7 × 20). In addition, we varied the hyperparameter search method (Optuna vs random search) and gene expression preprocessing (log_normalization vs expression binning with 20 bins). Fixed parameters included: XGBoost algorithm and balanced class weighting. Preprocessed data was generated once per percentile value and reused across all top_n_genes comparisons for that percentile.

Matthews Correlation Coefficient (MCC) was evaluated on the hold-out test dataset for each of the iterations, and values were plotted against number of retained features in the model (model size) either overall (Supplementary Figure 1A), grouped by percentile for clone summarization (Supplementary Figure 1B), by preprocessing method (Supplementary Figure 1C) or by hyperparameter tuning method (Supplementary Figure 1D). MCC is an appropriate metric for this evaluation as the positive class is represented at 10% of the total test set.

Average MCC increases as a function of top features retained from 10 through 50 and it plateaus out after 50. We selected 50 as the most parsimonious model size as it had higher MCC value (0.799) compared to 25 features (0.788, t-test p-value = 0.046), while there was no statistically significant difference between 50 and 75 (or higher) features.

Average MCC values trended higher for 75^th^ percentile for clone summarization, compared to 100^th^ and 50^th^ (median) percentiles, especially for smaller model sizes. 75^th^ percentile was chosen as default for training the model to account for any edge cases where 100^th^ percentile values are driven by outlier expression values in a cell within a clone.

Expression binning with 20 bins was overall superior to log normalization across multiple groupings (p-value < 0.0001). Expression binning with 10 bins was comparable to 20 bins for this dataset (not shown). Expression binning was also significantly faster (3-5x) to train a model compared to log normalization even when log normalized values were truncated to 4 decimals.

Finally, Optuna (Bayesian optimization) was superior to random search method for hyperparameter tuning across the different groups (p = 0.0049), and particularly for smaller model sizes.

##### Experiment 2: Algorithm Comparison

We evaluated three machine learning algorithms: XGBoost, AdaBoost, and Random Forest. The experiment compared 3 algorithms across 20 random seeds (seeds 1-20), resulting in 60 total training runs. Fixed parameters included: Optuna hyperparameter tuning, expression binning preprocessing with 20 bins, top 50 genes, and 75^th^ percentile for gene selection. All models used the same preprocessed data to enable direct comparison of algorithm performance.

##### Training Configuration

All experiments used nested cross-validation with 5 outer folds and 5 inner folds. Hyperparameter tuning was performed on the inner folds, and model evaluation was conducted on the outer folds. Cross-validation predictions were exported for downstream analysis. Feature selection was performed using highly variable gene selection with median aggregation across folds. All models were trained with class balancing enabled where applicable.

### 3 Data Availability

**Training datasets**

The Hanada *et al*., Lowery *et al.*, and Oliveira *et al.* datasets were downloaded from the dbGaP database.

The Hanada *et al*. dataset is available under accession phs002792.v1.p1: <https://www.ncbi.nlm.nih.gov/projects/gap/cgi-bin/study.cgi?study_id=phs002792.v1.p1>.

The Lowery *et al*. dataset is available under accession phs002748.v1.p1: <https://www.ncbi.nlm.nih.gov/projects/gap/cgi-bin/study.cgi?study_id=phs002748.v1.p1>.

The Oliveira *et al*. dataset is available under accession phs001415.v5.p1.c1: <https://www.ncbi.nlm.nih.gov/projects/gap/cgi-bin/study.cgi?study_id=phs001451.v5.p1>.

The Caushi *et al*. dataset was downloaded from the European Genome-Phenome archive, under study accession EGAS00001005343 (dataset EGAD00001007728): <https://ega-archive.org/datasets/EGAD00001007728>.

The Pétremand *et al.* dataset was downloaded from GEO and Zenodo: <https://www.ncbi.nlm.nih.gov/geo/query/acc.cgi?acc=GSE222448> and <https://doi.org/10.5281/zenodo.10869332>.

The Meng *et al.*, Gao *et al.*, and Ogura *et al.* datasets were downloaded from GEO.

The Meng *et al*. dataset is available at: <https://www.ncbi.nlm.nih.gov/geo/query/acc.cgi?acc=GSE254250>.

The Gao *et al.* dataset is available at: <https://www.ncbi.nlm.nih.gov/geo/query/acc.cgi?acc=GSE168859>.

The Ogura *et al.* dataset is available at <https://www.ncbi.nlm.nih.gov/geo/query/acc.cgi?acc=GSE209676>.

The 10x PBMC dataset was downloaded from the 10x Genomics site: <https://www.10xgenomics.com/datasets/human-pbmc-from-a-healthy-donor-10-k-cells-v-2-2-standard-5-0-0>.

**Atlas datasets**

The Zheng *et al.* pan-cancer TIL dataset was downloaded from Zenodo: <https://zenodo.org/records/5461803>.

The Liu *et al.* NSCLC dataset was downloaded from GEO: <https://www.ncbi.nlm.nih.gov/geo/query/acc.cgi?acc=GSE243013>.

The Salcher *et al.* NSCLC dataset was downloaded from Zenodo: <https://zenodo.org/records/7227571>.

The Chu *et al.* CRC dataset was downloaded from Figshare: <https://doi.org/10.6084/m9.figshare.25323397>.

The Loveless *et al.* PDAC datasets was downloaded from Zenodo: <https://zenodo.org/records/14199536>.

TRACE scores for the atlases shown in Figures 4 and 5 are available at <https://github.com/ksqtx/trace>.

### 4 Supplementary Figures and Tables

**Supplementary Table 2: Methodological comparison of published TRT prediction methods.** Details pertaining to published TRT methods and how some were applied to our aggregated dataset for benchmarking against TRACE.

| **Technique** | **Genes (method)** | **Scoring** | **Clone-level scores** | **Clone-level TRT calls** | **Notes (*)** |
| --- | --- | --- | --- | --- | --- |
| TRACE | 50 genes  (XGBoost tuning) | Individual cells or clones | Clone-level scores directly calculated | Threshold learned on training data, maximizing F1 |  |
| NeoTCR8 | 243 genes  (Seurat FindMarkers) | Individual cells (scGSEA) | Maximum cell score* | Threshold learned on training data, maximizing Youden’s J* | To apply NeoTCR8 to multiple training sets without clustering, the clone-wise score was set to the maximum cell score  A thresholding method was not identified, and Youden’s J was used. |
| TRTpred | 180 genes  (edgeR + QFL) | Individual cells (Singscore*) | Maximum cell score | Threshold learned on training data Z-scores, maximizing accuracy | The final scoring method was not identified, and Singscore was used. |
| TR30 | 30 genes  (Seurat FindMarkers) | Individual cells (UCell) | Mean cell score | Threshold learned on training data, maximizing Youden’s J* | A thresholding method was not identified, and Youden’s J was used. |
| MANAscore | 3 genes  (Authors’ selection) | Individual cells (non-imputed and imputed) | N/A; maximum cell score* | TRT clones are those containing ≥ 5 MANAscore^hi^ cells | The authors do not use a clone-wise score; for comparison, the maximum imputed MANAscore was used |
| predicTCR | >10 genes*  (XGBoost tuning) | Individual cells | Mean cell score | Fisher-Jenk break optimization | The full gene list was not provided and the code was unavailable |

**Supplementary Table 3: Overlap between genes used by TRACE and those used by other TRT prediction methods.**

| **Intersection** | **Intersection size** | **Genes** |
| --- | --- | --- |
| TRACE | 18 | *RBFOX2, LEF1, APOO, HLA-DQA1, CCL4L2, PRDX1, ID3, FASLG, RBM38, ANXA5, SLC9A9, PIK3R1, CREM, TLE5, NASP, FABP5, NR4A2, PDE4DIP* |
| TRACE & NeoTCR8 | 10 | *CCL4, HLA-DPA1, HLA-DRB1, CD63, PTTG1, ALOX5AP, CRIP1, CDC25B, JAML, LINC01871* |
| TRACE & TRTpred | 9 | *TNFRSF9, PHLDA1, KLF3, KLF2, GPR183, SELL, LTB, TCF7, DNAJB1* |
| TRACE & TR30 & TRTpred | 3 | *HAVCR2, CTLA4, VCAM1* |
| TRACE & TR30 & NeoTCR8 | 3 | *CXCR6, GZMB, NKG7* |
| TRACE & TRTpred & NeoTCR8 | 2 | *TOX, TIGIT* |
| TRACE & TR30 & TRTpred & NeoTCR8 | 2 | *DUSP4, LAG3* |
| TRACE & TRTpred & MANAscore | 1 | *IL7R* |
| TRACE & NeoTCR8 & MANAscore | 1 | *ENTPD1* |
| TRACE & TR30 & TRTpred & NeoTCR8 & MANAscore | 1 | *CXCL13* |

**Supplementary Table 4: Summary of TRT prediction method performance on the full and individual datasets**. Table shows test set MCC, F1, and AUC_PR scores for each model evaluated across 50 sets of holdout clones. Values are mean ± SD; *NA* MCC and F1 values were excluded from calculations.

| **Metric** | **Model** | **All datasets** | **Lowery *et al.*** | **Pétremand *et al*.** | **Meng *et al*.** | **Caushi *et al*. and Oliveira *et al.*** |
| --- | --- | --- | --- | --- | --- | --- |
| MCC | TRACE | 0.836 ± 0.032 | 0.804 ± 0.145 | 0.483 ± 0.256 | 0.440 ± 0.334 | 0.650 ± 0.097 |
|  | NeoTCR8 | 0.539 ± 0.032 | 0.766 ± 0.139 | 0.280 ± 0.291 | 0.089 ± 0.322 | 0.373 ± 0.129 |
|  | TRTpred | 0.806 ± 0.034 | 0.736 ± 0.144 | 0.570 ± 0.234 | 0.433 ± 0.224 | 0.574 ± 0.116 |
|  | TR30 | 0.592 ± 0.029 | 0.763 ± 0.147 | 0.574 ± 0.176 | 0.164 ± 0.458 | 0.433 ± 0.117 |
|  | MANAscore | 0.451 ± 0.066 | 0.310 ± 0.088 | 0.166 ± 0.121 | 0.041 ± 0.388 | 0.497 ± 0.124 |
| F1 | TRACE | 0.850 ± 0.029 | 0.880 ± 0.096 | 0.916 ± 0.047 | 0.862 ± 0.075 | 0.722 ± 0.086 |
|  | NeoTCR8 | 0.566 ± 0.027 | 0.868 ± 0.082 | 0.890 ± 0.059 | 0.814 ± 0.086 | 0.541 ± 0.100 |
|  | TRTpred | 0.821 ± 0.032 | 0.822 ± 0.103 | 0.925 ± 0.040 | 0.809 ± 0.086 | 0.667 ± 0.091 |
|  | TR30 | 0.606 ± 0.027 | 0.865 ± 0.088 | 0.931 ± 0.044 | 0.910 ± 0.070 | 0.580 ± 0.086 |
|  | MANAscore | 0.398 ± 0.067 | 0.295 ± 0.107 | 0.323 ± 0.117 | 0.477 ± 0.157 | 0.513 ± 0.129 |
| AUC_PR | TRACE | 0.890 ± 0.040 | 0.935 ± 0.086 | 0.934 ± 0.074 | 0.962 ± 0.041 | 0.763 ± 0.088 |
|  | NeoTCR8 | 0.677 ± 0.059 | 0.926 ± 0.058 | 0.928 ± 0.071 | 0.918 ± 0.082 | 0.636 ± 0.124 |
|  | TRTpred | 0.884 ± 0.032 | 0.955 ± 0.062 | 0.958 ± 0.061 | 0.953 ± 0.045 | 0.750 ± 0.095 |
|  | TR30 | 0.694 ± 0.053 | 0.886 ± 0.122 | 0.942 ± 0.067 | 0.940 ± 0.082 | 0.536 ± 0.135 |
|  | MANAscore | 0.817 ± 0.049 | 0.883 ± 0.094 | 0.936 ± 0.054 | 0.952 ± 0.059 | 0.660 ± 0.124 |

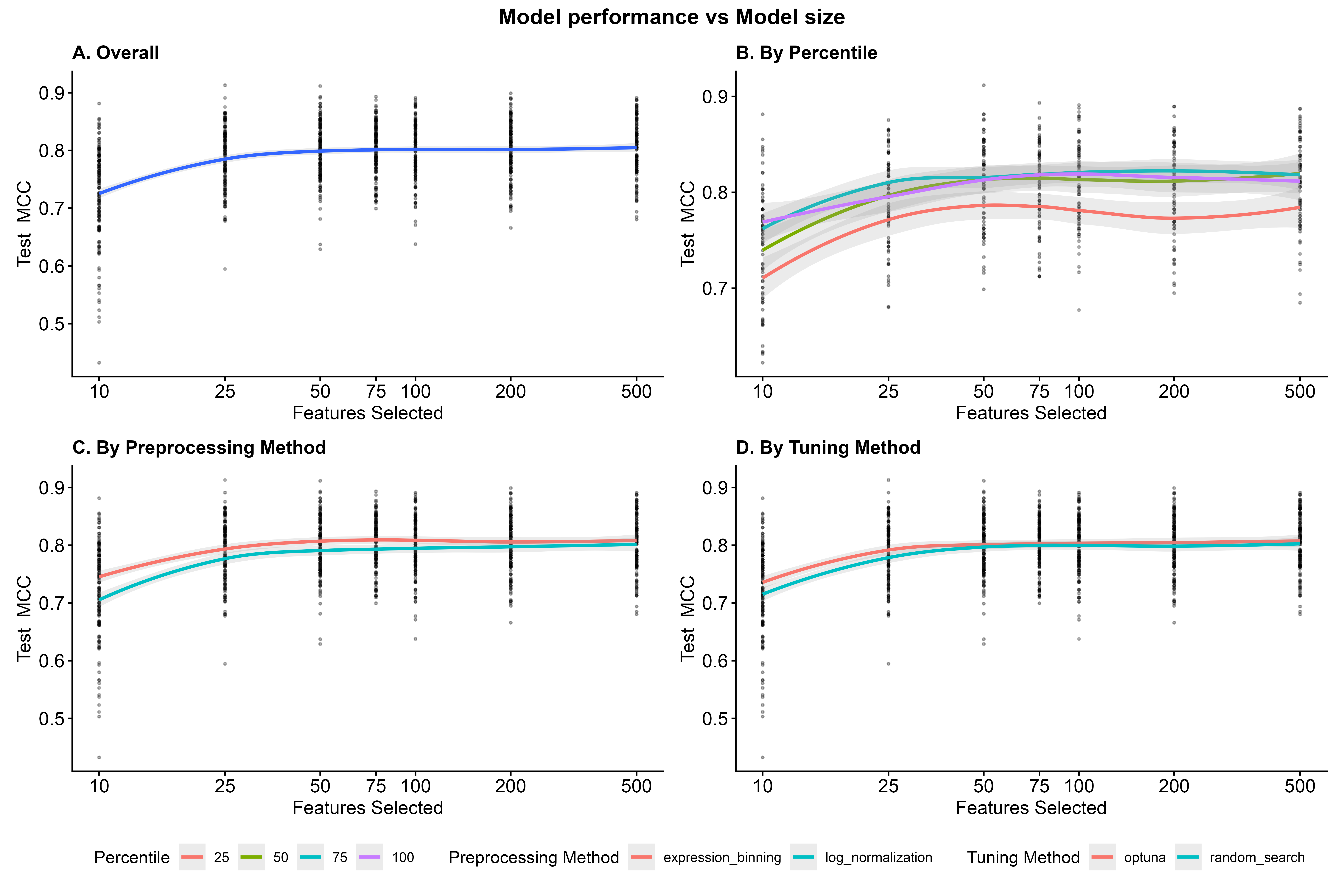

**Supplementary Figure 1.** **TRACE model performance in holdout test sets across 50 random iterations and range of tuning parameters.** **(A)** Test MCC increases with number of features selected in model up to 50 genes and incrementally beyond 50 genes. **(B)** Test MCC is grouped by gene expression percentile used for summarizing cells into clonal expression. **(C)** Test MCC is grouped by expression preprocessing method (expression binning in red; and log normalization in teal). **(D)** Test MCC is grouped by method of hyperparameter search (Optuna in red and Random grid search in teal).

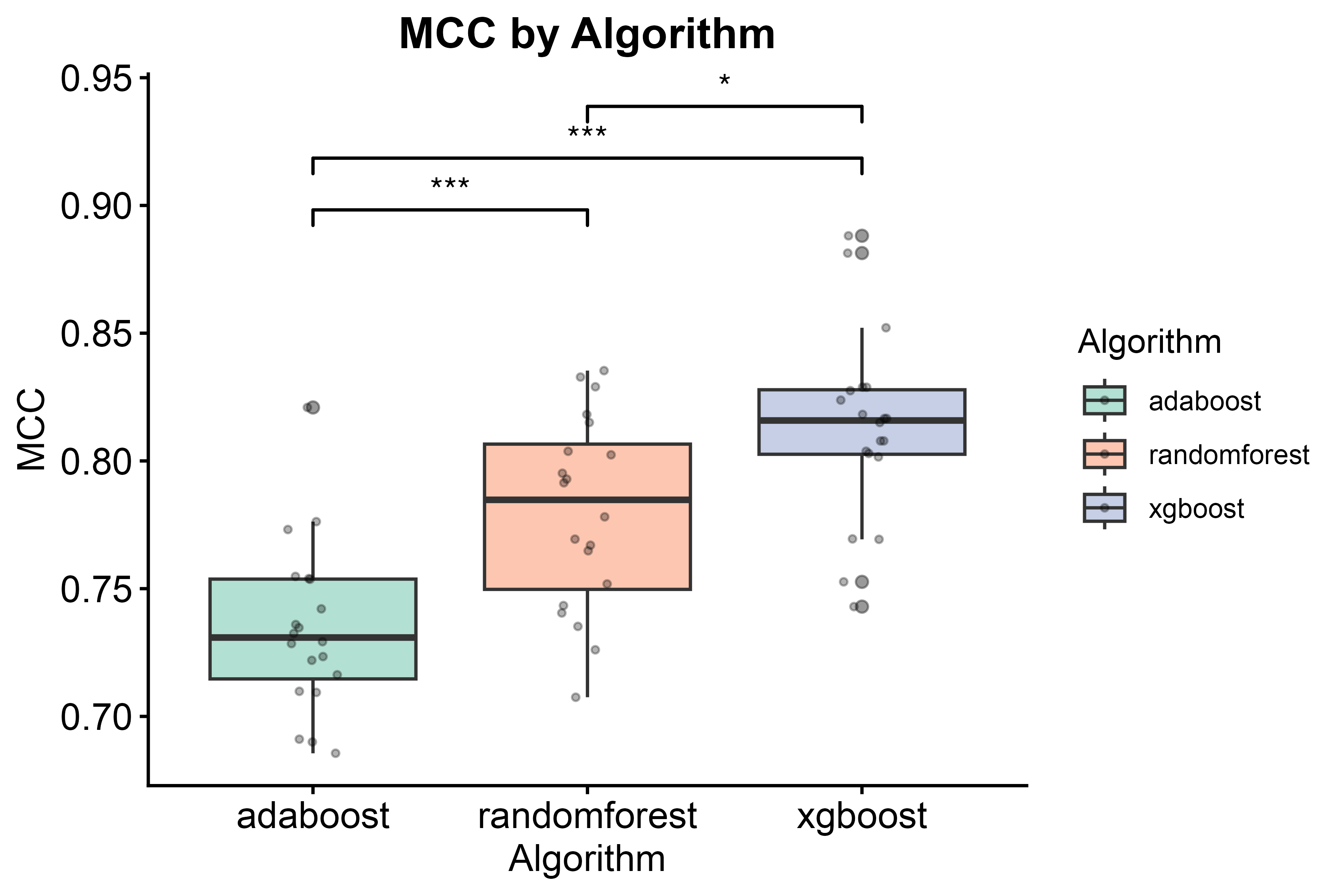

Supplementary Figure 2. Test MCC as a function of classification model. AdaBoost, RandomForest and XGBoost. (*p < 0.05; **p < 0.001)

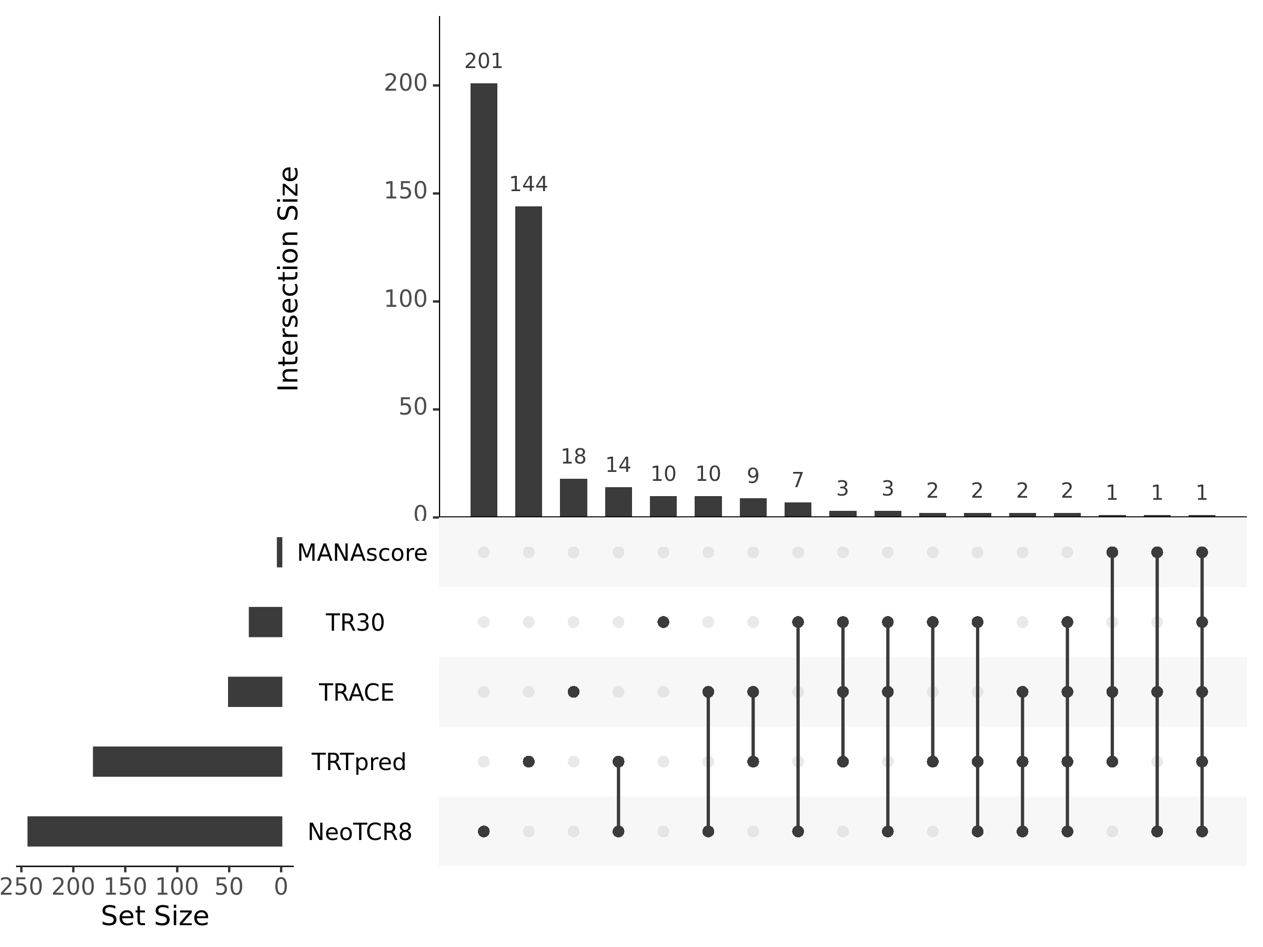

**Supplementary Figure 3. Sizes and intersections of feature sets used by TRACE and other TRT prediction methods used for benchmarking.** UpSet plot shows intersection sizes between TRACE genes and four other TRT methods.

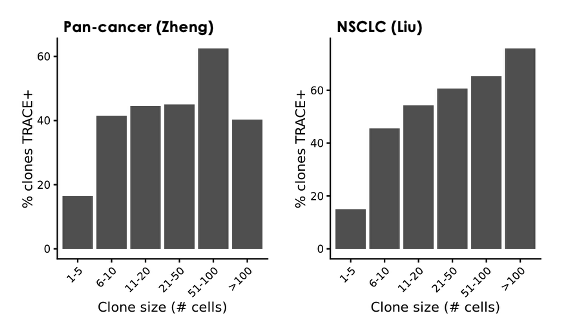

**Supplementary Figure 4. Percent of clones classified as TRACE^+^, binned by clone size.** Percentages are shown for the Zheng *et al*. pan-cancer TIL atlas (left) and the Liu *et al.* NSCLC study (right).

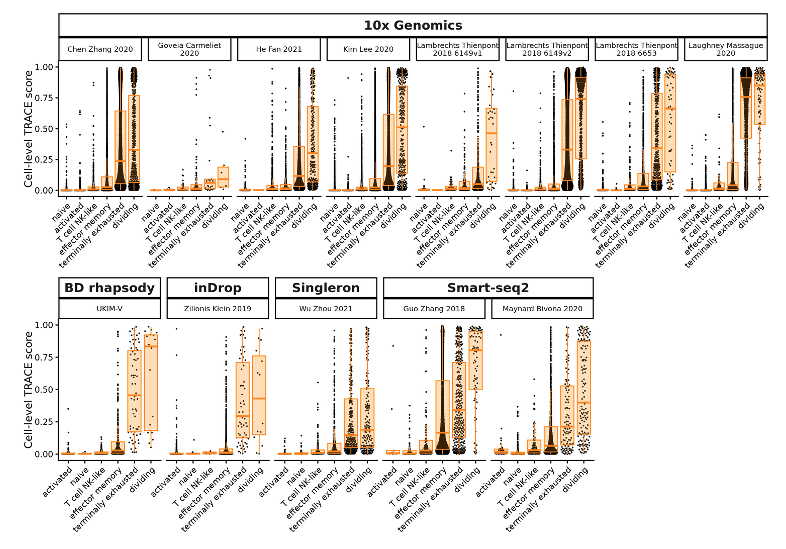

**Supplementary Figure 5. TRACE scores for Salcher *et al.* NSCLC atlas grouped by individual dataset and sequencing platform.** Scores are shown for tumor samples only and are grouped by CD8^+^ subtype.

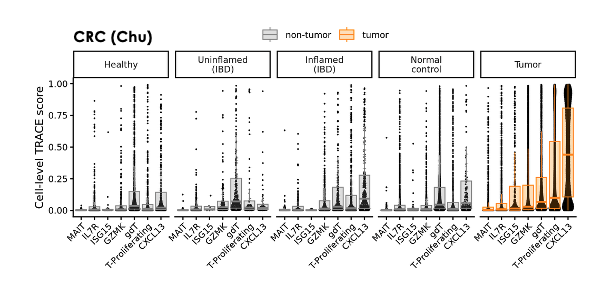

**Supplementary Figure 6. Distribution of TRACE scores across CD8^+^ T cells from the Chu *et al.* CRC atlas, grouped by CD8^+^ subtype.** Tumor cell scores are summarized with orange boxplots, and non-tumor cell scores are summarized with gray boxplots. Cells were subsampled down to 100 cells per sample to ensure relatively even representation of all samples.

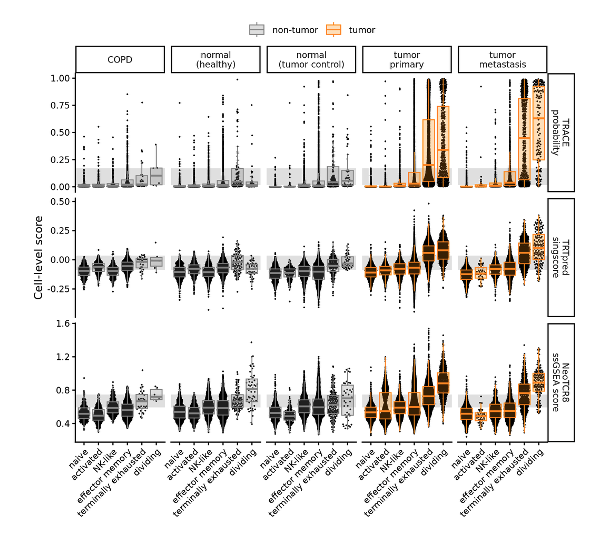

**Supplementary Figure 7. Comparison of tumor reactivity prediction methods using the Salcher *et al*. NSCLC atlas.** CD8^+^ T cells are grouped by CD8^+^ subtype and were subsampled down to 100 cells per sample to ensure relatively even representation of all samples. The same cells are shown for each prediction method. Tumor cell scores are summarized with orange boxplots, and non-tumor cell scores are summarized with gray boxplots. Gray shaded region depicts the upper and lower quartiles for terminally exhausted cells from normal (healthy) samples for each method.
